# Live imaging reveals chromatin compaction transitions and dynamic transcriptional bursting during stem cell differentiation *in vivo*

**DOI:** 10.1101/2022.10.07.511316

**Authors:** Dennis May, Sangwon Yun, David Gonzalez, Sangbum Park, Yanbo Chen, Elizabeth Lathrop, Biao Cai, Tianchi Xin, Hongyu Zhao, Siyuan Wang, Lauren E. Gonzalez, Katie Cockburn, Valentina Greco

## Abstract

Stem cell differentiation requires dramatic changes in gene expression and global remodeling of chromatin architecture. How and when chromatin remodels relative to the transcriptional, behavioral, and morphological changes during differentiation remain unclear, particularly in an intact tissue context. Here, we develop a quantitative pipeline which leverages fluorescently-tagged histones and longitudinal imaging to track large-scale chromatin compaction changes within individual cells in a live mouse. Applying this pipeline to epidermal stem cells, we reveal that cell-to-cell chromatin compaction heterogeneity within the stem cell compartment emerges independent of cell cycle status, and instead is reflective of differentiation status. Chromatin compaction state gradually transitions over days as differentiating cells exit the stem cell compartment. Moreover, establishing live imaging of *keratin-10* nascent RNA, which marks the onset of stem cell differentiation, we find that *keratin-10* transcription is highly dynamic and largely precedes the global chromatin compaction changes associated with differentiation. Together, these analyses reveal that stem cell differentiation involves dynamic transcriptional states and gradual chromatin rearrangement.

## Introduction

Cellular identity is a composite of many features, including behavior, morphology, protein levels, and gene expression. All of these aspects are fundamentally shaped by the transcriptional program and therefore chromatin architecture of a cell. Recent technological advances have allowed the field to increasingly appreciate transcriptional heterogeneity within cell populations that were previously assumed to be homogeneous (Patel et al., 2014; Wang et al., 2020), and single-cell epigenetic profiling is beginning to reveal the extent of chromatin architecture heterogeneity within cell populations (Buenrostro et al., 2015; Finn et al., 2019; Jin et al., 2015; Lai et al., 2018)

The particular suite of genes expressed in a given cell is largely determined by nucleosome compaction in different region of the genome, where accessible (euchromatic) regions permit gene expression, and inaccessible (heterochromatic) regions largely prevent gene expression. During embryonic lineage specification, stem cell differentiation, and somatic cell reprogramming, chromatin architecture undergoes large-scale changes resulting in drastically different cell identities (Golkaram et al., 2017; Kurimoto et al., 2015; Oudelaar et al., 2020; Paulsen et al., 2019; Pelham-Webb et al., 2020). However, because chromatin architecture analyses, including those with single-cell resolution, typically rely on data captured at fixed timepoints, we lack an understanding of how chromatin architecture progressively changes during cell identity transitions within a physiological setting.

Epidermal stem cell differentiation is an excellent model to understand the progressive nature of cell identity transitions. The epidermis is fueled by a basal layer of stem cells which are actively proliferating to maintain density, and the continual differentiation of cells delaminating and moving apically to build the outer layers of the skin barrier (**Supp Fig 1a**). Traditionally, cell identities within the epidermis have been distinguished by cell morphology, specific protein markers, and localization within the tissue.

Recently, single cell RNA-sequencing data has shown that the cell identity transition through epidermal differentiation is progressive and takes place over several days. Specifically, cells within the basal stem cell layer show global transcriptional changes associated with differentiation preceding exit from the basal layer (Aragona et al., 2020; Cockburn et al., 2021). Chromatin accessibility and the architecture of individual loci have been investigated in embryonic skin (Fan et al., 2018; Gdula et al., 2013; Mardaryev et al., 2014; Shue et al., 2020), but how and when chromatin changes relative to the transcriptional and morphological transitions of adult epidermal stem cell differentiation remains unknown.

Here, we leverage intravital imaging to observe and track global chromatin changes of individual stem cells within their homeostatic environment through time and cell fate transitions. By developing a quantitative pipeline to capture each cell’s unique chromatin compaction state, we reveal extensive heterogeneity of global chromatin architecture within the epidermal stem cell population, as well as distinct chromatin compaction states of epidermal stem cells and their differentiated daughter cells. Tracking individual cells over time and using a reporter for differentiation status reveals that global chromatin compaction state reflects differentiation state, beginning in the basal layer prior to exit from the stem cell compartment. We also show, through live imaging endogenous transcription at the earliest known stage of differentiation, that epidermal cells pass through heterogeneous and flexible transcriptional states as they progress towards their fully differentiated status. Together, this study reveals the chromatin compaction heterogeneity within a regenerative organ, incremental chromatin compaction remodeling through stem cell differentiation, and insight into how transcriptional dynamics of a key differentiation gene relate to cellular state transitions.

## Results

### Intravital imaging reveals cell cycle-independent heterogeneity in chromatin compaction across the basal stem cell layer

To understand transitions in large-scale chromatin architecture as a function of cell identity, we developed a fluorescence-based system that allows visualization of chromatin compaction in single skin epidermal stem cells within their native tissue in live mice. This quantitative pipeline leverages fluorescently-tagged histone 2b where bright fluorescence indicates densely packed chromatin and dimmer fluorescence indicates loosely packed chromatin or chromatin-excluded compartments (Amiad-Pavlov et al., 2021; Kanda et al., 1998). Using the *keratin-14 histone2B-GFP* (*K14H2B-GFP*) allele, which is expressed in epidermal stem cells and their progeny, we first segmented the 3D volume of individual nuclei and extracted fluorescence intensity at each voxel from high-resolution, intravital imaging data (**Fig 1A and B Movie 1**). This complex, spatial dataset was then reduced to a unique intensity distribution profile by normalizing all voxels within the 3D volume and plotting them as a percentage of total nuclear volume (**Fig 1C**). The normalization also accounts for any variation in raw intensity values among different nuclei, allowing for comparison among different cell populations and mice, irrespective of differences in mean intensity. Therefore, any changes in chromatin compaction profiles reflect the relative change in chromatin architecture intrinsic to the individual nucleus, that can therefore be compared to any other analyzed nuclei.

**Figure 1:**
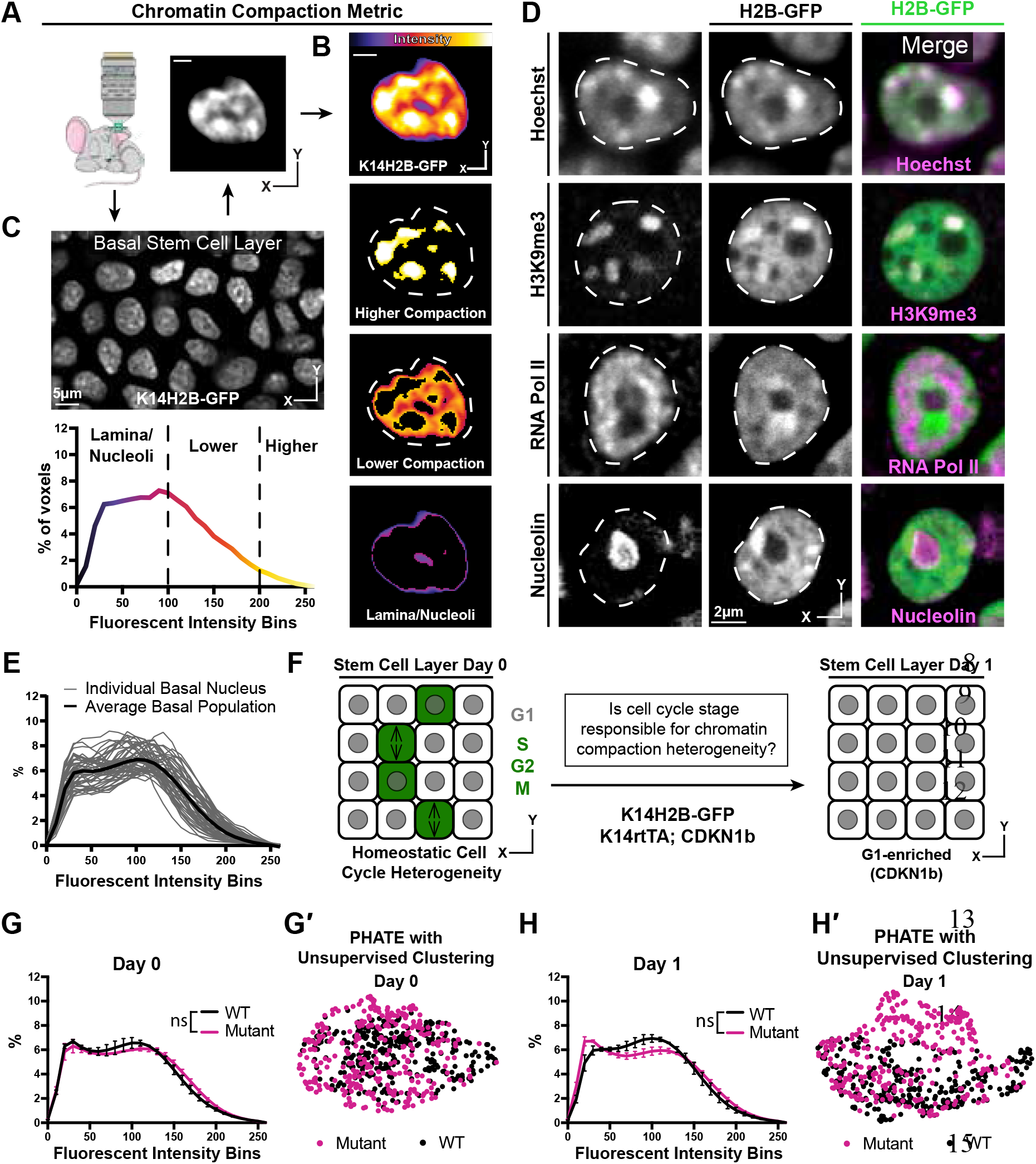
Chromatin compaction state is heterogeneous and independent of interphase cell cycle. **(A)** Representative XY view of the basal stem cell layer showing the *kertain14*-driven Histone2B-GFP allele in a live mouse. **(B)** A representative chromatin compaction profile of a single basal stem cell nucleus. Each voxel from the 3D volume of a nucleus was exported, normalized for mean fluorescent intensity, and plotted as a voxel percentage of volume against the 0-256 intensity bins (methods). **(C)** A representative nucleus expressing H2B-GFP in a single optical slice, where the fluorescence intensity is displayed as a heatmap to illustrate the range of chromatin compaction within a nucleus. See Movie 1 for a 3D rendering of this nucleus. **(D)** Fixed epidermal tissue expressing H2B-GFP (green) and co-stained with Hoechst or various subnuclear compartment markers (magenta), demonstrating that high H2B-GFP fluorescence intensity correlates with heterochromatin (H3K9me3), lower H2B-GFP fluorescence intensity correlates with euchromatin (RNA Pol II), and the lowest H2B-GFP fluorescence intensity (in part) correlates with nucleoli (Nucleolin). The colocalization/overlap of these two fluorophores are indicated by the presence of white signal. Nuclear outlines are traced in white, dotted lines. **(E)** Chromatin compaction plots for 50 individual basal stem cells (grey lines) and the averaged population (bold, black line) revealing substantial heterogeneity of chromatin compaction states within the stem cell population. **(F)** Schematic of the genetic p27 (CDKN1b) overexpression system to stall cells in late G1 after 1 day of doxycycline administration. An increased N of 1260 basal nuclei across all mice (3 mutant and 3 wild-type) over day 0 and day 1 because only a subset of basal stem cells are within a non-G1 cell cycle phase at any given time. **(G)** Comparison of chromatin compaction states of wild-type (K14H2B-GFP; K14rtTA) and mutant (K14H2B-GFP; K14rtTA; tetO-CDKN1b) mice prior to doxycycline administration/induction on day 0. **(G′)** PHATE plot of the same wild-type and mutant cells from panel (G) on day 0 showing intermixed populations. **(H)** Comparison of chromatin compaction states of wild-type (K14H2B-GFP; K14rtTA) and mutant (K14H2B-GFP; K14rtTA; tetO-CDKN1b) mice one day post doxycycline induction and stalling of the cell cycle in late G1 showing non-significant changes between wild-type and mutant populations. **(H′)** PHATE plot of the same wild-type and mutant cells from panel (H) on day 1 after CDKN1b induction showing largely intermixed populations.

Applying this pipeline to intravital imaging data allowed us to quantify chromatin compaction within individual nuclei, among a population of stem cells, and between distinct cell identities based on cell location within the skin of a live mouse (**Supp fig 1B and 1C**).

The variable intensity of H2B-GFP indicates regions with different levels of chromatin compaction: highly-compressed, constitutively repressed chromocenters, loosely packed euchromatin, and nuclear periphery and nucleoli (**Fig 1C**). In fixed tissue staining, we recapitulated these chromatin compaction regions with Hoechst, as well as localization of known subnuclear compartments such as H3K9me3-positive chromocenters in the high H2B-GFP intensity regions, active RNA polymerase in the lower H2B-GFP intensity regions, and nucleoli in the very low H2B-GFP intensity regions (**Fig 1D, Supp fig 2A**). Applying the histone deacetylase inhibitor Trichostatin-A (TSA) abrogated these differences in H2B-GFP fluorescent intensity throughout individual nuclei and resulted in a shifted chromatin compaction curve compared to that of epidermal cells in the stem cell layer treated with the DMSO vehicle control (**Supp fig 2B and 2D**). Thus, the relative H2B-GFP intensity provides a visual readout of large-scale chromatin architecture (**Fig 1C**) and can be quantified to visualize the relative distribution of chromatin at different levels of compaction within individual nuclei irrespective of their volumes and mean H2B-GFP fluorescence intensities (**Fig 1B**).

Applying our chromatin compaction analysis to the basal stem cell layer, we noticed a clear heterogeneity within the stem cell population (**Fig 1E**). Previous studies have demonstrated chromatin organization heterogeneity at the level of individual locus accessibility or TAD boundaries (Buenrostro et al., 2015; Finn et al., 2019; Jin et al., 2015; Lai et al., 2018), but given that our chromatin compaction analysis captures very large-scale aspects of global chromatin architecture, the degree of heterogeneity we observed in the basal stem cell layer was surprising. Cells within the stem cell layer are in a spread of cell cycle states at any given time, so we hypothesized that cells may differentially condense their chromatin throughout the interphase cell cycle, producing an overall heterogeneity in chromatin compaction states reflective of cell cycle status. At any given point, ∼20% of the cells within the basal layer are in S/G2/M, and the remaining ∼80% are in G1 (Hiratsuka et al., 2015). By inducing overexpression of the cell cycle inhibitor p27 (CDKN1b), we stalled the entire basal stem cell population in G1 (*K14rtTA; tetO-CDKN1b; K14H2B-GFP*) (**Fig 1F**). Intriguingly, reducing cell cycle heterogeneity did not significantly change the population-level chromatin compaction state of basal stem cells (**Fig 1G, 1H**). To explore more subtle relationships between individual cell’s chromatin compactions states, we used the dimensionality-reducing data visualization algorithm, PHATE (Moon et al., 2019). In this method, individual data points represent single cells, and the distance between points reflects the similarity of those cells’ chromatin compaction profiles. In our data, the proportion of each nucleus’ H2B-GFP intensity in each bin is analogous to each cell’s number of reads for each gene in the more typical single cell RNA-sequencing application of PHATE (**Supp fig 2C**). Plotting the combined wild-type and mutant populations together revealed that they largely intermix both within the homeostatic cell cycle distribution (day 0) and through the cell cycle stall in G1 (day 1) (**Fig 1G′ and 1H′)**, which is in contrast to the largely separated populations of cells treated with TSA vs. DMSO (**Supp fig 2D’**). These data are consistent with previous studies which showed heterogeneity in chromatin architecture and accessibility independent of cell cycle differences (Buenrostro et al., 2015, 2015; Finn et al., 2019; Lai et al., 2018). Together, these results show that single cell chromatin compaction states in the basal stem cell layer are heterogeneous and independent of interphase cell cycle status.

### Differentiation status of epidermal cells dictates chromatin compaction state

The epidermis is a highly-regenerative organ involving the continual differentiation of cells from the basal stem cell layer to form the functional barrier of our skin. Cells within the stem cell layer will continually alter their transcriptome, delaminate from the stem cell compartment, and move apically/outward to form the overlaying differentiated layer (called spinous) over the course of 3-4 days (Cockburn et al., 2021; Mesa et al., 2018) (**Fig 2A**). Previous studies have demonstrated that basal stem cells and differentiated spinous cells have different nuclear volumes and numbers of pericentromeric clusters (Gdula et al., 2013). With live imaging of H2B-GFP, we observe additional qualitative differences in chromatin architecture between these two populations; for example, differentiated spinous cells have a higher number of compact, bright chromocenters, and overall flatter nuclei (**Fig 2B**). By applying our chromatin compaction analysis to these two different cell populations, we observed that they have distinct chromatin compaction profiles, where the compaction state of differentiated cells was shifted toward lower fluorescent intensity bins (**Fig 2C**). As the quantitative pipeline allows for comparison between nuclei irrespective of differences in mean fluorescent intensity values, the shift left for differentiated cells implies a relative increase in very low-intensity regions such as the nuclear lamina and nucleoli, with a relative decrease in euchromatic and heterochromatin regions (**Supp fig 1B**).

**Figure 2:**
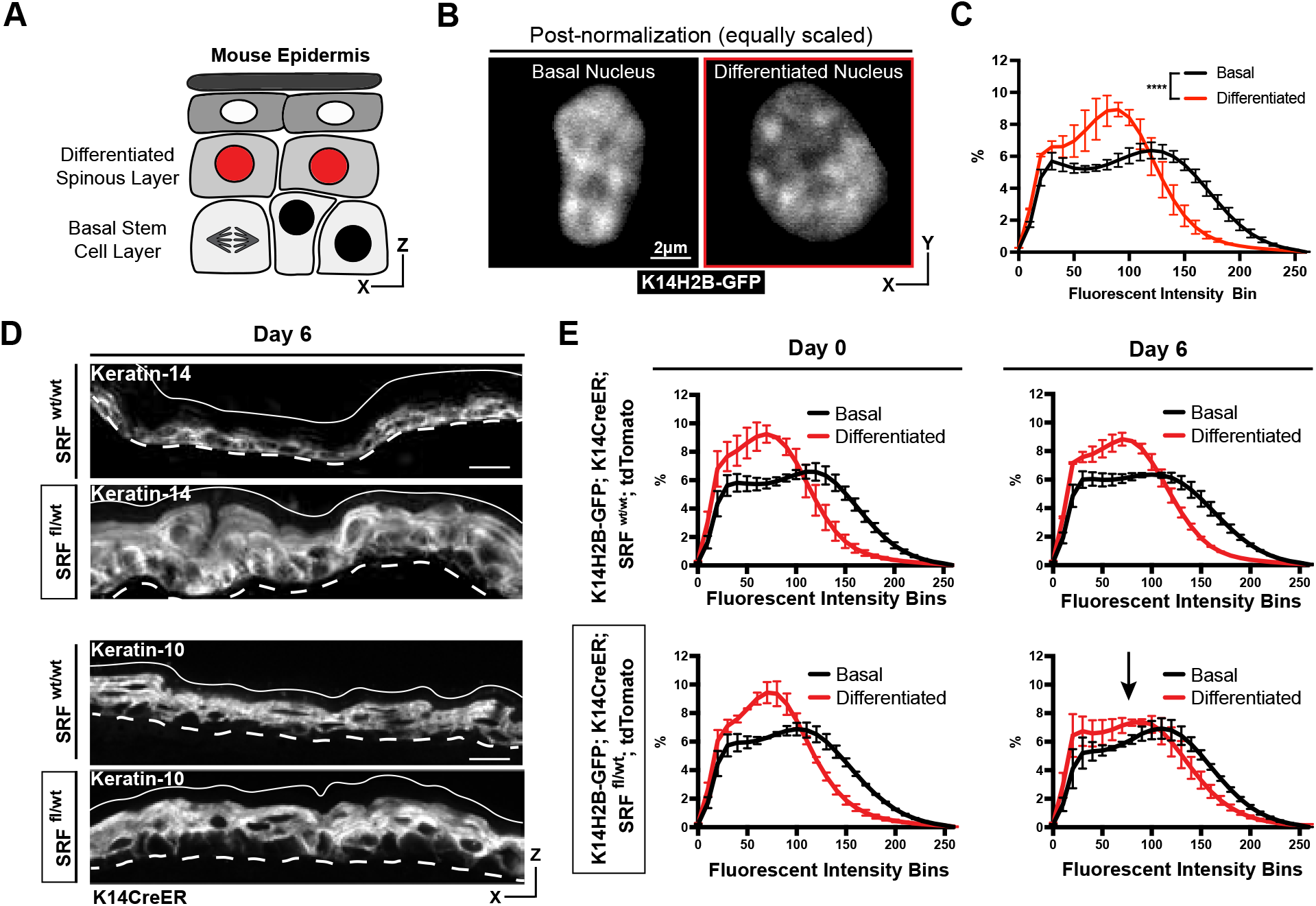
Chromatin compaction state changes through differentiation state. **(A)** XZ schematic of the epidermis. The basal stem cell layer is shown with black nuclei and the differentiated (spinous) layer shown apical with red nuclei. **(B)** Representative crops of individual nuclei from the basal and differentiated populations scaled identically in the first two panels, and scaled individually in the third to better show the different chromatin compaction states between cell identities. H2B-GFP signal in white. **(C)** Chromatin compaction profiles of averaged populations of cells from the basal stem cell and differentiated layers showing significant differences in chromatin plots. N = 150 basal and 90 spinous cells across 3 mice. **(D)** Fixed, XZ tissue slices from SRF^wt/wt^ and SRF^fl/WT^ mice day 6 after tamoxifen recombination. The basal stem cell marker, Keratin-14, can be seen expanded into differentiated layers, and the differentiated marker, Keratin-10, can be seen localized correctly despite a thickened overall epidermis. **(E)** Chromatin compaction profiles for wild-type (SRF^wt/wt^; K14CreER; K14H2B-GFP; tdTomato) and mutant (SRF^fl/wt^; K14CreER; K14H2B-GFP; tdTomato) mice on day 0 and 6 after tamoxifen recombination. Black arrow in SRF^fl/wt^ day 6 denotes that significant change in spinous cell chromatin compaction profile. N = 150 basal and 90 spinous cells across 3 mice per day.

To better understand how differentiation affects chromatin compaction state, we genetically knocked out one copy of *Serum Response Factor* (SRF) in the epidermis of adult mice (*K14H2B-GFP; SRF*^*fl+*^; *K14CreER; LSL-tdTomato*) (**Supp fig 3A**). SRF is a transcription factor which helps establish proper cell identity in the epidermis, contributing both to embryonic skin stratification and proper stem cell differentiation in the adult mouse epidermis (Lin et al., 2013; Verdoni et al., 2010). At 6 days post-recombination, the loss of SRF caused a transcriptional identity shift in the skin; expression of the basal stem cell marker Keratin-14 was no longer restricted to the basal stem cell layer, despite grossly normal epidermis organization (**Fig 2D, Supp fig 3B**). In both SRF heterozygous and wild type control mice, the chromatin compaction states in basal and spinous cells were distinct on day 0. By day 6, the differentiated, spinous population’s chromatin compaction was more basal like in the SRF heterozygous mice, while the WT controls maintained distinct chromatin compaction profiles between the cell identities (**Fig 2E**). In particular, the chromatin compaction state of spinous differentiated cells shifted towards the chromatin compaction state of basal stem cells, reflecting the expansion of Keratin-14 expression into the spinous layer and the breakdown of transcriptome identity between the two cell populations. Importantly, these data also demonstrate that chromatin compaction changes can be reflective of changes in transcriptional program and cell identity, even prior to extensive tissue phenotypes.

### Basal cell chromatin compaction is stable over hours, but transitions through differentiation over days

Because we observed that chromatin is differentially compacted and organized between the basal stem cell and differentiated (spinous) layer above, we hypothesized that chromatin architecture would reorganize at a specific transition point during stem cell differentiation. To test this hypothesis, we began by analyzing chromatin compaction from three-hour timelapse imaging of *K14H2B-GFP* mice. Visually, chromatin compaction in cells in the basal stem cell layer was relatively stable over this time. High H2B-GFP density regions appeared to move only slightly, and chromatin compaction profiles at the beginning, middle, and end of these three-hour time lapses exhibited no significant change at the individual cell or population level (**Fig 3A and A′, Supp fig 4A**). These findings demonstrated that among these heterogeneous basal stem cells, the chromatin compaction state of each cell is stable over hours.

**Figure 3:**
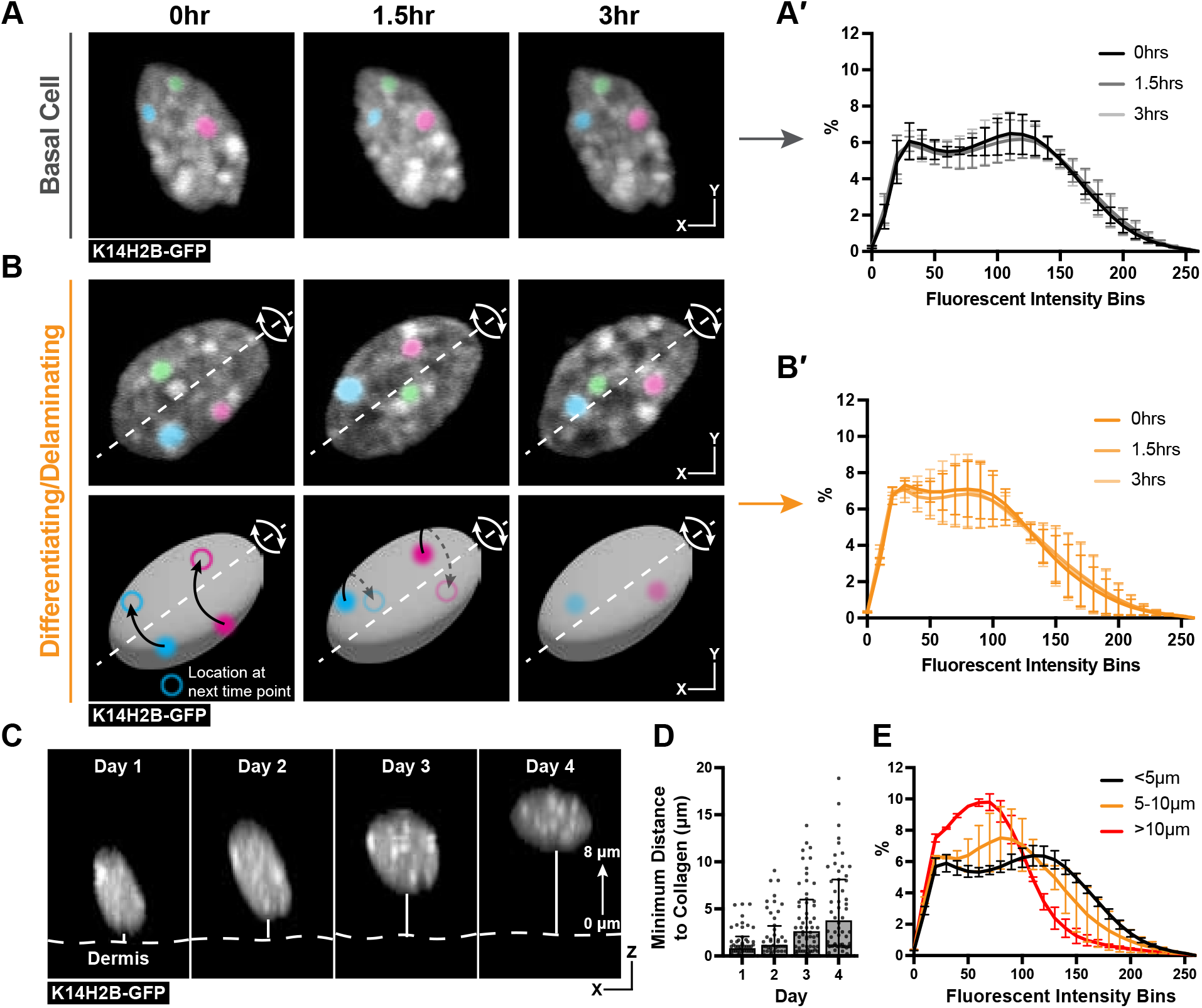
Chromatin compaction is stable over hours and progressively changes over days. **(A)** Time lapse imaging data of a single nucleus crop at 0, 1.5, and 3-hour time points. H2B-GFP fluorescent signal is shown in white. Three, high-intensity chromocenters were chosen and pseudo-colored blue, green, and pink to demonstrate their static nature over the 3 hours. **(A′)** Chromatin compaction profiles for cells in the basal stem cell layer at the 0, 1.5, and 3-hour time points showing no significant change over 3 hours. N = 150 basal cells across 3 mice. **(B)** Same as in (A) but cells actively delaminating out of the basal stem cell layer, exhibiting rolling nuclei. The axis of rotation is show in the white dotted line, with the rotational direction shown in white arrows around that axis. Three, high-intensity chromocenters were chosen and pseudo-colored blue, green, and pink to demonstrate the dynamic spinning taking place, but the positional stability of global chromatin organization relative to itself. A cartoon (below) of the same nucleus to better visualize the rotation and orientation over the 3 hours with the blue and pink pseudocolored chromocenters tracked through time. Black arrows indicate where the chromocenter will move to in the next time point (hollow circle), dotted black arrow indicates rotation around the backside of the nucleus. **(B′)** Chromatin compaction profiles for cells with spinning chromatin (actively delaminating cells) showing no significant change over 3 hours. N = 146 spinning/delaminating cells over 3 mice. **(C)** XZ crops of the same nucleus tracked within the tissue over 4 days. Representative example of a differentiating cell over this time period. Bold, dotted white line denotes the epidermal/dermal. Solid, thin white line shows the minimum distance from collagen quantified in panel (D). H2B-GFP fluorescent signal shown in white. **(D)** Minimum distance from collagen for a randomly selected population of basal cells on day 1, some of which differentiated and moved apically by day 4, while others remained basally located. **(E)** Chromatin compaction profiles of nuclei within the tracked population binned as distance from collagen showing the direct transition in chromatin compaction profiles through differentiation. N = 150 basal stem cells tracked over 4 days from 3 mice.

We next wondered whether the transition in chromatin compaction occurs slightly later during differentiation in cells actively delaminating, located in between the basal and spinous layer (Cockburn et al., 2021) (**Supp Fig 4B**). Strikingly, we noticed that the chromatin of these cells spins over the course of the timelapse (**Fig 3B, Movie 2**). The chromatin spinning was observed in all timelapses obtained under homeostatic conditions. Surprisingly, even in such a dramatic and dynamic nucleus, chromatin compaction was maintained through spinning with no significant changes in chromatin compaction taking place (**Fig 3B′**). Notably, though, the chromatin compaction state of these actively delaminating cells was markedly different from the chromatin compaction state of basal cells (compare **3B′** to **3A′**), and appeared to be an intermediate differentiation state between basal and spinous cells. Overall, these timelapse data suggest the transition in chromatin compaction state during epidermal differentiation occurs over days, not hours, and is already in progress during basal cell delamination.

To test this hypothesis, we tracked a population of cells in the basal stem cell layer over four days in a live mouse (**Fig 3C**). By doing so, we sought to understand when and how quickly chromatin compaction changes were taking place, as well as confirming that the cells progressively transition between the chromatin states seen in Fig 2B and 2C. This population increased their average minimum distance from collagen from day 1 to day 4, reflecting that a portion of these cells were differentiating and moving apically into the spinous layer of the epidermis (**Fig 3C and 3D, Supp fig 5A**). After tracking all cells over this time period, we binned nuclei into groups determined by their minimum distance to collagen to quantitatively measure the delamination process. When the chromatin compaction analysis was applied to these populations, we observed a gradual and directional change in chromatin compaction state as cells moved farther from collagen and moved into the spinus layer (**Fig 3E**). Together, our results indicate that chromatin compaction remodels slowly over days concomitant with differentiation and delamination.

### Chromatin compaction state begins to transition within the basal stem cell layer as differentiation initiates

The observation that delaminating basal cells had chromatin compaction states similar to differentiated spinous cells (**Fig 3B-B′**) made us wonder whether chromatin reorganization began while cells were still within the basal stem cell layer. This model could also explain the heterogeneity of chromatin compaction states within the basal layer (**Fig 1E**). Indeed, previous studies have shown that basal stem cell differentiation involves cumulative transcriptional changes that begin prior to delamination (Aragona et al., 2020; Cockburn et al., 2021). We wondered if the chromatin compaction state of basal cells that are committed to differentiation differs from basal cells that are not.

To test this hypothesis, we genetically labelled the cells of the basal stem cell layer with an early marker of differentiation: expression of *Keratin-10* (*K10rtTA; tetO-Cre; tdTomato; K14H2B-GFP*) (Cockburn et al., 2021; Muroyama and Lechler, 2017) (**Fig 4A and 4B, Supp fig 5B**). This population represents a relatively large portion of the basal stem cell layer (∼40%), and expression of the K10 reporter has been shown to define a point of commitment for delamination (Cockburn et al., 2021). Basal cells were sampled with respect to K10 reporter (tdTomato) expression, and then binned into K10+ and K10-groups. Remarkably, even this relatively small difference in cell states (all a part of the basal stem cell layer) had significantly different chromatin compaction profiles (**Fig 4C**). In addition, the cells positive for the K10 reporter adopted an intermediate chromatin compaction profile which was shifted towards that of the fully differentiated cells in Fig 2C and delaminating/differentiating cells in Fig 3B^**′**^. These data highlight that basal stem cells initiate global chromatin changes coinciding with delamination and exit from the stem cell compartment.

**Figure 4:**
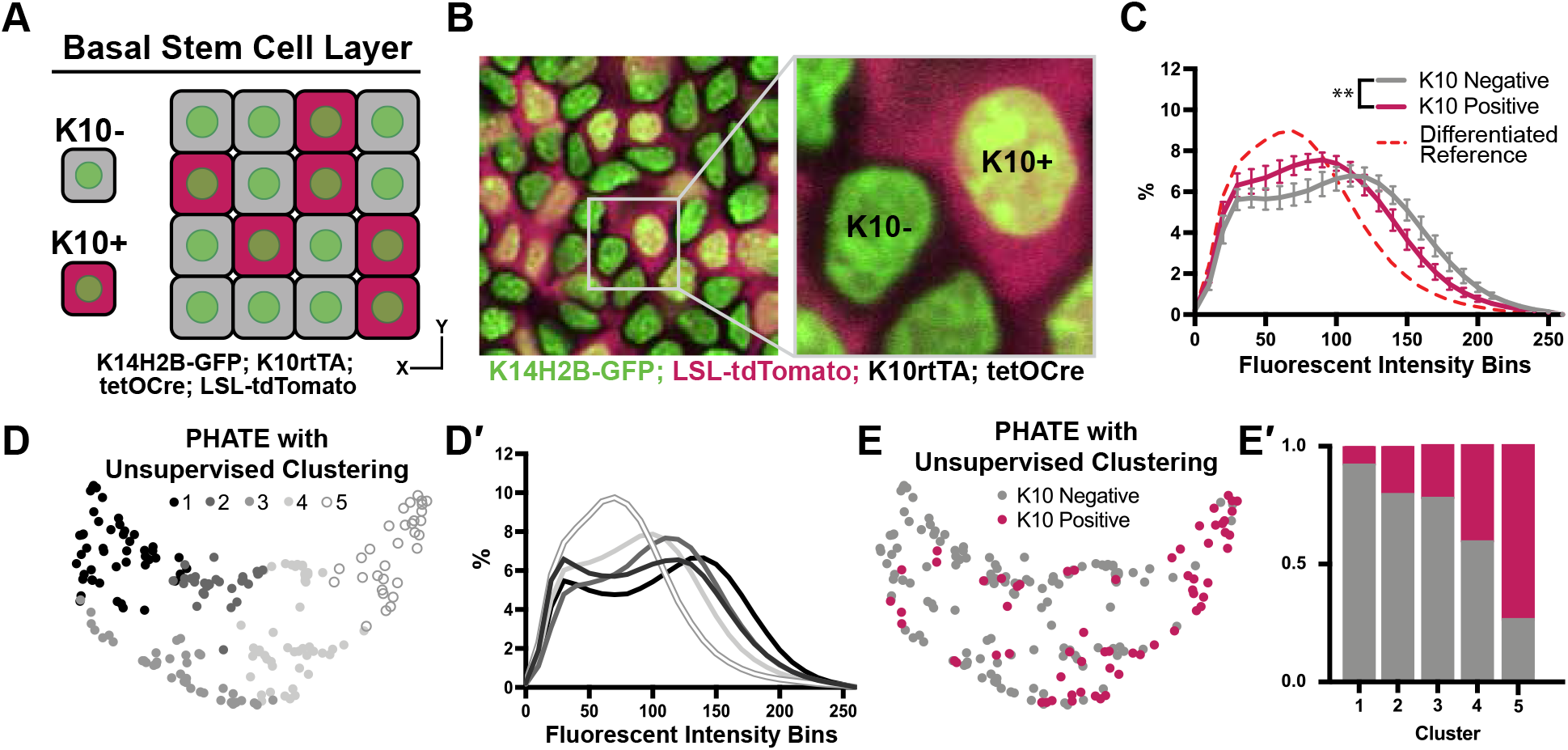
Chromatin compaction changes precede differentiation. **(A)** XY schematic of genetic system (K14H2B-GFP; K10rtTA; tetO-Cre; tdTomato) allowing visualization of actively differentiating cells still within the basal stem cell layer (expressing differentiation-associated *keratin-10* gene). *Keratin-10* positive cells indicated with red cytosol. **(B)** Representative XY crop of the basal stem cell layer showing *keratin-10* negative (no tdTomato signal) and *keratin-10* positive cells (tdTomato in cytosol) with a cropped inset on the right. **(C)** Chromatin compaction profiles comparing K10 status (tdTomato on/off) in basal stem cells showing significant differences in chromatin compaction between groups. A differentiated reference line is shown by the dotted, red curve. Averaged K10+ cells shown in pink and averaged K10-cells shown in grey. N=187 K10-nuclei and N=132 K10+ nuclei from 3 mice. Statistical comparisons made between histogram groups, p<0.01. **(D)** PHATE plot of data from (C). Louvain clustering results are projected onto the PHATE plot. Each dot represents one nucleus profile, and the distance between dots represents the similarity in chromatin compaction profile. **(D′)** Chromatin compaction profiles of the averaged clusters identified through Louvain clustering in (D) elucidating directionality in the clustering from a more basal curve to a more differentiated curve. **(E)** The same PHATE/clustering dataset as in (D) with overlayed K10 status (on/off) again demonstrating directionality in the PHATE map. **(E′)** The ratio of K10 positive (red) and negative (grey) in each of the Louvain clusters.

Applying the PHATE analysis and an unsupervised Louvain clustering algorithm to this basal stem cell dataset (including both K10- and K10+ cells), five distinct clusters emerged (**Fig 4D**). The chromatin compaction profiles of the cells in the five clusters showed an entire spread of curves from most “ basal-like” (cluster 1) to nearly fully differentiated (cluster 5) (**Fig 4D**^**′**^), reflecting the overall differentiation trajectory of chromatin compaction states. Together with the shape of the PHATE map itself, we noticed that the left and right sides (clusters 1 and 5) of the map seemed to narrow and pinch together, implying more similar chromatin compaction states among cells at the beginning and end of this trajectory than among the cells in the middle of the trajectory (clusters 2-4) (**Fig 4D)**. Because these maps were derived from the K10 reporter dataset, we were able to overlay each cell’s K10 status onto the PHATE maps (**Fig 4E**). The distribution of K10+ and K10-cells within the PHATE map clusters (**Fig 4E**^**′**^) reinforced the directionality of chromatin compaction changes indicated in Fig 4D′. Together, these data show that changes in chromatin compaction coincides with stem cell commitment to differentiation.

### Dynamic Keratin-10 expression precedes global chromatin compaction changes in differentiating basal stem cells

Intrigued by how the differentiation trajectory seemed to be reflected in chromatin compaction states, we wanted to more specifically understand the relationship between global chromatin compaction state and transcription at the *keratin-10* locus. To do so, we used the MS2/MCP genetic system to visualize *keratin-10* transcripts in real time. We knocked in the MS2 cassette (24x repeats of the MS2 stem loop sequence) after the stop codon of the *keratin-10* locus, and then crossed this mouse line to the MCP-GFP reporter mouse (Lionnet et al., 2011) and TIGRE-K14-H2B-mCherry (methods). MCP-GFP binds the MS2 stem loops, and this happens as soon as *keratin-10-MS* is transcribed. Thus, these mice (*Keratin-10MS2/+; MCP-GFP/+; TIGRE-K14-H2B-mCherry*) allowed us to visualize not only chromatin architecture through mCherry-tagged H2B, but also *in vivo* transcription of an endogenous allele through a GFP-positive punctum in the nucleus (**Fig 5A**). Importantly, while both the *Keratin-10-MS2/MCP-GFP* transcription reporter and the K10rtTA reporter in Fig 4 will capture cells that have been stably expressing *keratin-10* for some time, the transcription reporter can capture cells at an earlier stage of *keratin-10* expression compared to the K10rtTA reporter, which relies on multiple successive steps of gene expression, translation, and recombination.

**Figure 5:**
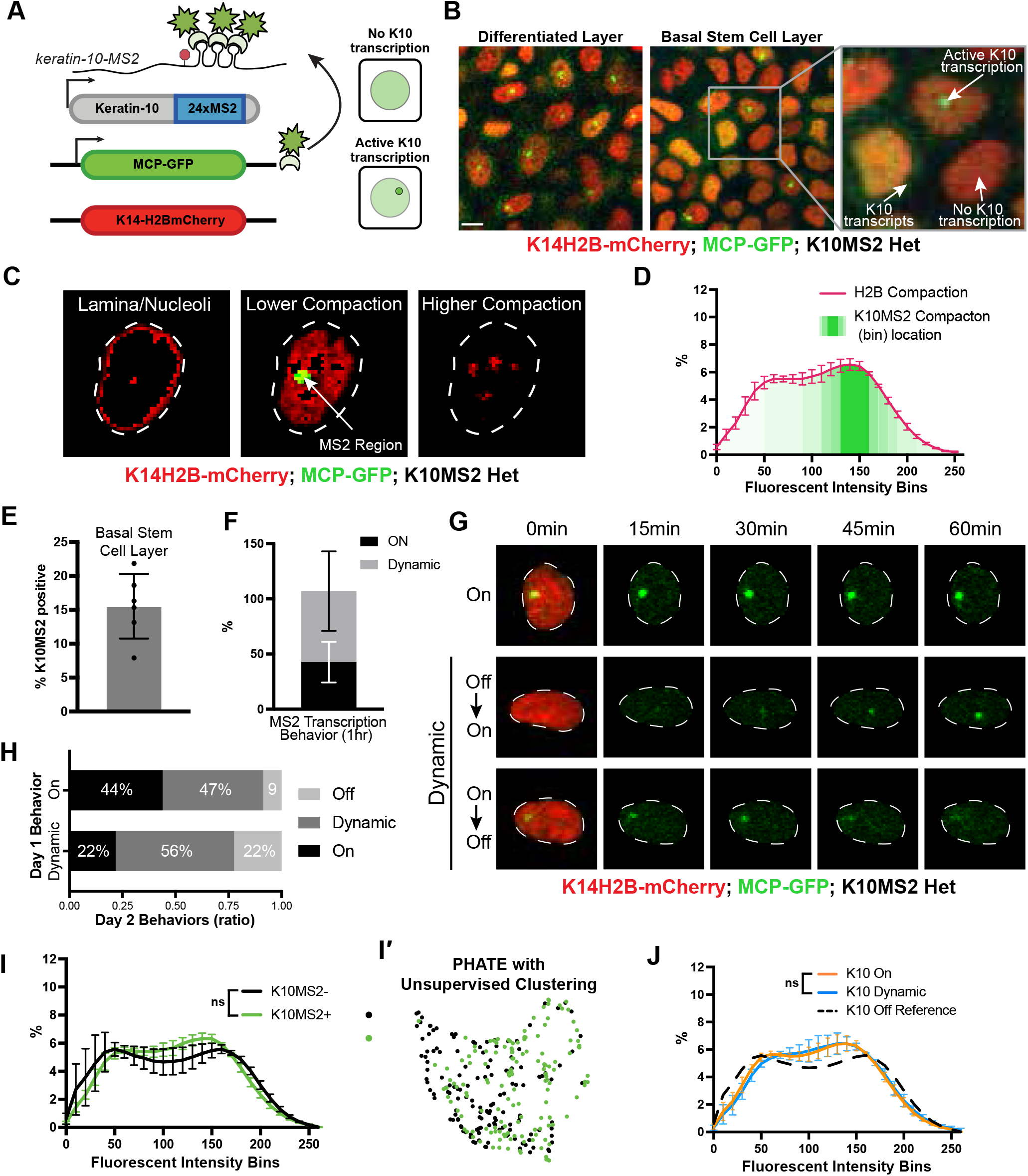
*in vivo* transcription of *keratin-10* precedes genome architecture changes through differentiation. **(A)** Visual schematic of the MCP/MS2 system allowing visualization of a targeted gene under endogenous regulation. 24X MS2 repeats were knocked into the 3’UTR of the *keratin-10* locus and chromatin compaction visualized with K14H2B-mCherry. Presence of nuclear MCP punctum indicates active *keratin-10* transcription, where lack of signal indicates no active transcription at that locus. **(B)** Representative crops of the differentiated spinous layer (left) and basal stem cell layer (right), as well as magnified inset showing active site of *keratin-10* transcription, and nascent transcripts in cytosol awaiting translation. Scale bar=5μm. **(C)** Representative image of a nucleus separated into the same chromatin compaction regions as in Fig 1C. Active transcription of *keratin-10* can be seen within the lower compaction region from bins 100-200. **(D)** Chromatin compaction profile of averaged H2B-mCherry basal stem cells (red line) and the fluorescent bin location of *keratin-10* transcription punctum (green, semi-translucent bars). N = 150 *keratin-10* transcribing basal stem cells across 3 mice. Surfaced MCP signal was used to identify the mCherry fluorescent intensity bins that *keratin-10* transcription occurred, and then plotted as increasing green transparency. **(E)** Populational percentage of active *keratin-10* transcription in the basal stem cell layer in K10MS2 Het mice. N = 3 100 × 100um regions quantified over 3 mice. **(F)** Percentages of *keratin-10* dynamics within the *kertain-10* positive basal stem cell layer over 1 hour. ∼40% of basal stem cells actively expressing *keratin-10* remained on throughout the 1-hour timelapse, while ∼60% had dynamic transcriptional behaviors. **(G)** Representative crops of individual basal stem cell nuclei exhibiting different *keratin-10* transcriptional dynamics over the course of 1 hour. H2B-mCherry signal can be seen in red in the first time point for reference, and *keratin-10* transcription punctum in green in all time points. White, dotted line shows the nuclear border. **(H)** Quantification of how *keratin-10* dynamics change over the course of 1 day. *Keratin-10* positive nuclei from day 0 were binned into “ on” and “ dynamic” (y-axis) and the same nuclei located on day 1. Day 1 transcriptional dynamics were quantified (x-axis) showing surprising flexibility in transcriptional dynamics over a day. **(I)** Chromatin compaction analysis of basal stem cells either actively transcribing *keratin-10* (green punctum in nucleus) or not transcribing *keratin-10*. Despite active transcription of the differentiation gene, there is no significant chromatin compaction remodeling at this stage. N = 150 *keratin-10* “ off” and 150 “ on” nuclei across 3 mice. **(I′)** PHATE and Louvain clustering of the data in (I) showing intermixed populations of *keratin-10* positive and negative basal stem cells. **(J)** Chromatin compaction analysis of the “ on” and “ dynamic” *keratin-10* transcription populations from (F) showing very little, non-significant differences between the two populations. The *keratin-10* “ off” curve from (I) is shown as a reference in the dotted, black line N = 187 “ dynamic” and 124 “ on” over 3 mice.

Imaging the epidermis of these mice confirmed that differentiated spinous cells had active transcription of *keratin-10*, and only a subset of cells in the basal stem cell layer had active *keratin-10* transcription at any given time (**Fig 5B, 5E**), which is consistent with previous characterizations of Keratin-10 protein expression patterns (Braun et al., 2003; Doupé et al., 2010; Schweizer et al., 1984). To further validate that MCP-GFP puncta represent active *keratin-10* transcription at its locus, we next sought to resolve the local chromatin environment at the *keratin-10* locus during active transcription. To do so, all H2B-mCherry fluorescence voxels within the nuclear surface of MCP-GFP puncta-containing basal stem cells were normalized as previously described, and those voxels that overlapped with the surface of MCP-GFP signal were extracted, enabling a quantitative visualization of the H2B-mCherry fluorescence within the local chromatin environment of *keratin-10* transcription. This analysis revealed that *keratin-10* transcription primarily occurred within loosely packed euchromatic regions of the nucleus (**Fig 5C and 5D, Supp fig 5D**), as expected. Interestingly, cells in the basal layer displayed three different reporter localization patterns: no transcription of *keratin-10* (no green punctum in nucleus), active transcription of *keratin-10* (bright punctum in nucleus), and multiple, dimmer MCP-GFP puncta in the cytosol without active transcription of *keratin-10* (no nuclear punctum) (**Fig 5B**). This last group led us to believe transcription of *keratin-10* might be quite dynamic, with kinetics in the range of the half-life of the *keratin-10* mRNA itself, likely minutes to hours (Dar et al., 2012; Suter et al., 2011). To understand possible *keratin-10* transcription dynamics, we imaged *Keratin10-MS2/+; MCP-GFP/+; K14H2B-mCherry* mice over the course of one hour. Intriguingly, of the cells actively transcribing *keratin-10*, about 40% sustain *keratin-10* transcription with no notable change in fluorescence intensity, but about 60% displayed dynamic transcription over the timelapse – they turned *keratin-10* on or turned *keratin-10* off during the hour imaged (**Fig 5F and 5G**). This pattern is consistent with most genes being transcribed in “ bursts” lasting less than one hour (Bothma et al., 2014; Chubb et al., 2006; Dar et al., 2012; Fritzsch et al., 2018; Suter et al., 2011). To determine the perdurance of transcriptional dynamics, we performed timelapse imaging of the same exact cells 24 hours later. We observed that only a subset of cells displayed the same transcriptional behaviors between the two days – approximately 40% of *keratin-10* “ on” cells were “ on” the next day, while roughly 60% *keratin-10* “ dynamic” cells were “ dynamic” the next day. The remaining cells which had been transcribing *keratin-10* on day 1 displayed a variety of different *keratin-10* transcriptional behaviors on day 2, including an absence of active transcription entirely (**Fig 5H**). Collectively, these data indicate that *keratin-10* transcriptional activity in basal stem cells is flexibly dynamic.

To understand how *keratin-10* transcriptional status relates to chromatin compaction states, we used the H2B-mCherry signal to perform chromatin compaction analysis. This analysis revealed that cells actively transcribing *keratin-10* had chromatin compaction states similar to that of cells not transcribing *keratin-10* (**Fig 5I and 5I′, Supp fig 5C**), though they trended towards the chromatin compaction state of cells expressing the K10rtTA reporter (compare to **Fig 5I** to **Fig 4C**). Because the cells marked by the *Keratin-10-MS2/MCP-GFP* transcription reporter include those at the very earliest step of differentiation, this result suggests that chromatin compaction changes either have not yet occurred or are just beginning to occur when *keratin-10* transcription is initiated.

Finally, we hypothesized that *keratin-10* “ on” cells were further advanced in their cell identity transition toward differentiation than *keratin-10* “ dynamic” ones. Intriguingly, the chromatin compaction signature of *keratin-10* “ on” compared to *keratin-10* “ dynamic” cells were extremely close to one another (**Fig 5J**) suggesting that the cells displaying these different *keratin-10* transcriptional dynamics in fact co-exist within extremely similar cell identity states.

Altogether, these results reveal that the initiation of *keratin-10* transcription, which represents one of the first steps of a basal stem cell towards differentiation, is highly dynamic and that significant chromatin remodeling occurs after transcription initiation of *keratin-10*.

## Discussion

Homeostasis and function of regenerative tissues requires constant self-renewal and differentiation of resident stem cells. Stem cells undergo a relatively large reorganization of their genome through the differentiation process as their cell identity changes (Kurimoto et al., 2015; Li et al., 2017; Oudelaar et al., 2020; Paulsen et al., 2019). Current tools offer high resolution data into the chromatin environment at specific loci, but rely on fixed cells and are unable to reveal how chromatin architecture is remodeled during cell identity changes such as differentiation. Here, through high-resolution imaging of H2B-GFP in live mice, we discovered chromatin compaction heterogeneity within the epidermal stem cell population under homeostatic equilibrium. This heterogeneity arises from gradual, incremental shifts in stem cell identity throughout differentiation which begin prior to exit from the stem cell layer. Moreover, by live imaging endogenous transcription of *keratin-10*, a hallmark of epidermal differentiation, we resolved highly dynamic transcriptional activity over hours and days in the absence of significant chromatin architecture changes. Ultimately, we determined that most of the global genome reorganization associated with cell identity transitions occurs between the initiation of differentiation-associated transcription and exit from the stem cell layer (delamination) (**Figure 6, model**).

**Figure 6:**
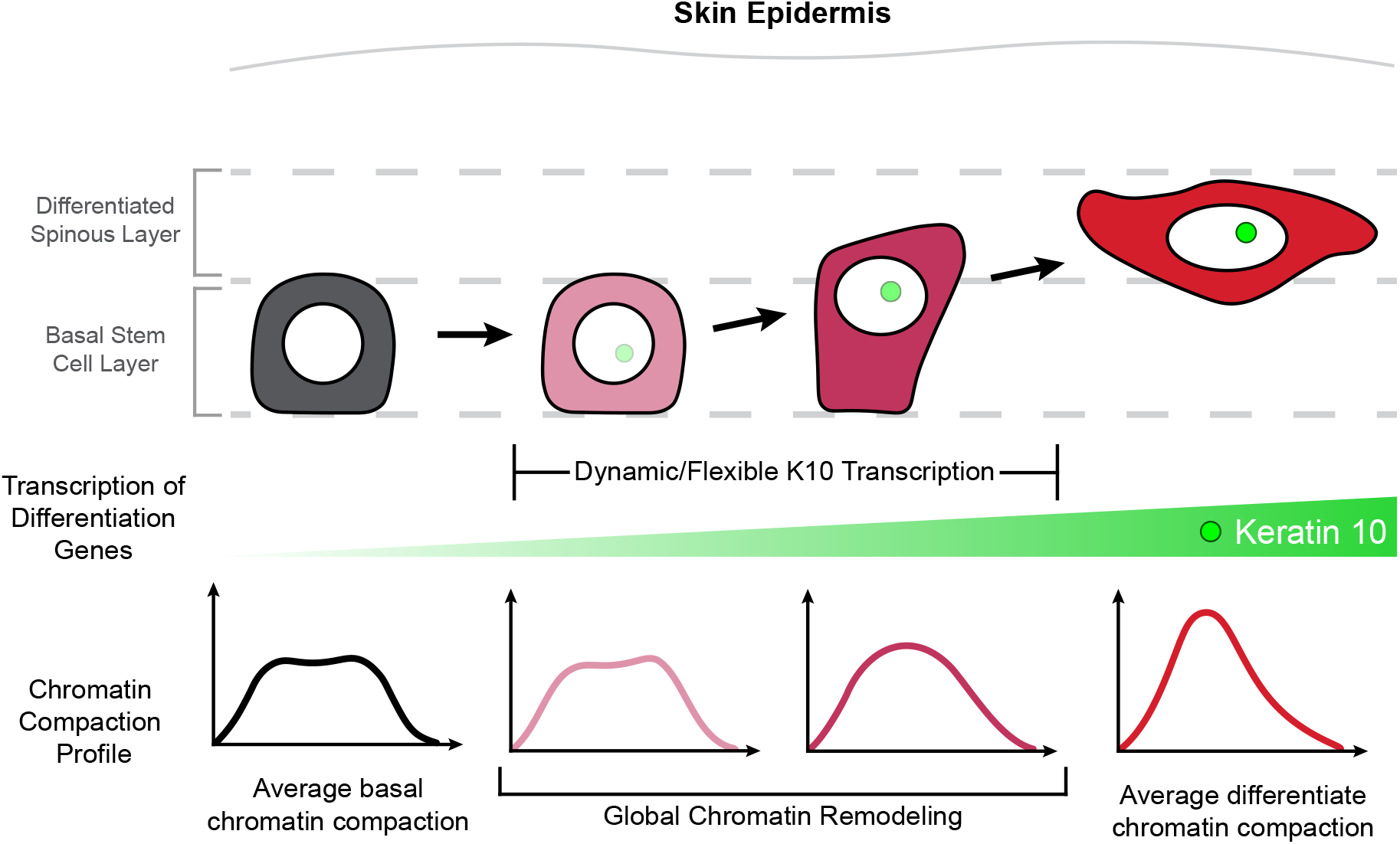
Chromatin architecture remodeling through epidermal differentiation. Epidermal stem cells undergo incremental changes toward differentiation over 3-4 days (top). During this process, differentiation-committed cells within the basal stem cell layer begin expressing *keratin-10* dynamically over hours and flexibly over days (green circle/site of transcription in nucleus). Basal stem cell and fully differentiated spinous cells have specific chromatin architecture states associated with their different cellular identities (bottom row), and global chromatin remodeling begins before delamination and exit from the basal stem cell layer.

Our ability to live image chromatin compaction in the same cells over hours and days allowed us to discover that the global chromatin compaction state of epidermal stem cells is stable over the course of hours, but remodels through the differentiation process over days. Recently, other studies which performed live imaging in cell culture systems to evaluate chromatin dynamics at finer scales, such as individual loci and Topologically Associated Domain (TAD) boundaries, have revealed that chromatin can be locally dynamic over minutes and seconds (Barth et al., 2020; Chen et al., 2018; Gabriele et al., 2022; Iida et al., 2022). These dynamic local changes to individual loci and TADs may act cumulatively toward a global and incremental shift in chromatin architecture through differentiation.

By visualizing chromatin compaction changes at the same time as morphological and transcriptional changes, our results add to a growing understanding that the first steps of epidermal stem cell differentiation are somewhat flexible. Recent work tracking epidermal cell fates over a week showed that differentiating basal cells expressing *keratin-10* are still capable of proliferation (Cockburn et al., 2021), a behavior that was previously assumed to be unique to epidermal stem cells. Additionally, pseudo-time analysis of scRNA-sequencing data from the same study revealed a significant population of cells that contain both stem and differentiation-associated transcripts (*keratin-14* and *keratin-10*, respectively). While the dynamic nature of *keratin-10* transcription we observed within a one-hour timelapse could reflect the fact that that many genes exhibit transcriptional bursting kinetics on the scale of minutes (Dar et al., 2012; Suter et al., 2011), the observation that a portion of cells actively transcribing *keratin-10* no longer express it one day later supports the idea that individual cells remain somewhat flexible in their initial commitment to differentiation. Understanding at what point individual cells irreversibly commit to differentiation and can no longer proliferate, as well as if chromatin compaction state is also flexible during differentiation, remain interesting questions for future studies.

Finally, we have begun to tease apart the intimate relationship between chromatin architecture changes and transcriptional behavior *in vivo*. By combining two different genetic approaches to temporally visualize differentiation state (**Fig 4** and **Fig 5**), our results suggest that transcriptional changes happen either before or right at the beginning of global chromatin remodeling. Recent papers have supported the bi-directional and reciprocal nature of chromatin organization at individual gene loci and transcription of those loci (Lai et al., 2018; Li et al., 2021; Oudelaar et al., 2020). Through our live imaging approach, we have greatly enriched our understanding of the interplay between the highly dynamic nature of transcription and local chromatin architecture by tracking cell identity transitions in a live mammal.

Finally, this chromatin compaction system is not dependent on biology inherent to the skin or epidermis, and in fact is widely applicable to other systems due to the use of H2B-GFP in many model systems. Additionally, this approach could be used to investigate other biological transitions in cell identity such as oncogenic initiation and expansion, and mesenchymal transitions through wound healing. There is also an opportunity to combine this more global view of chromatin architecture with higher resolution imaging modalities of individual loci or TAD boundaries. More broadly, these findings open a door into tissue-level coordination and flexibility among cells, and how the incremental and stepwise journey through differentiation establish heterogeneous cell states. We believe taking a more global view of these individual cell states, such as this tracking of pan-histone labeling, is one avenue to understand such processes.

## Author contributions

DM, SY, LG, KC, and VG designed experiments and wrote the manuscript. DM, SY performed experiments. DM, SY, and DG analyzed data. DM and DG developed chromatin compaction profiling pipeline. EL performed SRF tissue sectioning. YC and SW performed PHATE and Louvain clustering analysis. BC and HZ developed statistical analysis to compare chromatin compaction histograms. SP characterized chromatin spinning phenotype after discovery.

## Acknowledgements

We thank all members of the Greco lab for critical feedback on the manuscript. We thank T. Lechler for K10-rtTA mice, E. Fuchs for K14H2B-GFP mice, K. Politi for tetO-Cre mice, as well as Vladimir Botchkarev (Boston University), Tatiana Efimova (George Washington University), Joerg Bewersdorf (Yale University), Jennifer Philips-Cremins (University of Pennsylvania), Shangqin Guo (Yale Univeristy), and Maria Kasper and Karl Annusver (Karolinska Institute) for their kind feedback at various stages of the project. We also thank the Yale Transgenic Facility for their work in generating the TIGRE-H2B-mCherry and K10MS2 mouse lines, and the Yale intravital imaging core for their continued support.

This work is supported by an HHMI Scholar award and NIH grants number 1R01AR063663-01, 1R01AR067755-01A1, 1DP1AG066590-01 and R01AR072668 (VG). SY is supported by a Medical Scientist Training Program grant from the NIH (T32GM136651), and DM is supported by the NIH-funded Training Program in Genetics (T32GM007223-44) and the Lo Stem Cell Fellowship (Yale).

## Methods

### *In vivo* imaging

All imaging was performed in non-cycling regions of the ear skin with hair removed using depilatory cream (Nair) before the start of each experiment. Mice were anesthetized using 1-2% vaporized isoflurane delivered by a nose cone throughout the course of imaging. Image stacks were acquired with a LaVision TriM Scope II (LaVision Biotec, Germany) laser scanning microscope equipped with both a Chameleon Vision II and Discovery 2-photon lasers (Coherent, USA). For collection of serial optical sections, the laser beam was focused through a 40x water immersion lens (Nikon; N.A. 1.15) and scanned with a field of view of 200×200um at 600Hz. Z-stacks were acquired with 0.5-1μm steps to image a total depth of ∼40μm of tissue, covering the entire thickness of the epidermis. Visualization of ECM was achieved via second harmonic signal using blue channel at 940 nm imaging wavelength. To follow the same epidermal cells over multiple days, inherent landmarks of the skin together with a micro-tattoo were used to navigate back to the same epidermal regions every 24h. For time-lapse imaging, serial optical sections were obtained in a range of 15-30 minute intervals for a total duration of 1-3h.

### Immunofluorescence

For SRF tissue-section analysis, ear skin was dissected, fixed with 4% paraformaldehyde in PBS for 1hr at room temperature and then embedded in optimal cutting temperature (OCT; Tissue Tek). Frozen OCT blocks were sectioned at 10μm. Primary antibodies used were guinea pig anti-K10 (1:200; Progen PG-K10) and rabbit anti-K14 (1:200; BioLegend 905301). All secondary antibodies used were raised in a donkey host and were conjugated to AlexaFluor 568 or 633 (Thermofisher). Fixed tissue was mounted on a slide with Vectashield Anti-fade mounting medium (Vector Laboratories) with a #1.5 coverslip.

To isolate epidermis for whole mount staining, ear tissue was incubated in 5mg/ml dispase II solution (Sigma, 4942078001) at 37°C for 10 minutes and the epidermis was separated from dermis using forceps. Epidermal tissue was fixed in 4% paraformaldehyde in PBS for 45 minutes at room temperature, washed 3X in PBS, permeabilized and blocked for >1h (2% Triton-X, 5% Normal Donkey Serum, 1% BSA in PBS), incubated in primary antibody overnight at 4°C, and secondary antibodies for 3h at room temperature the next morning. Primary antibodies used were as follows: guinea pig anti-K10 (1:200; Progen GP-K10), rabbit anti-K14 (1:200; BioLegend 905301) rabbit anti-H3K9me3 (1:200; Abcam ab8898), rabbit anti-nucleolin (1:200; Abcam ab22758), and rabbit anti RNA Polymerase pS2 (1:500; Abcam ab5095). All secondary antibodies used were raised in a donkey host and were conjugated to AlexaFluor 568 or 633 (Thermofisher). Fixed tissue was mounted on a slide with Vectashield Anti-fade mounting medium (Vector Laboratories) with a #1.5 coverslip.

### Image analysis

Raw image stacks were imported into FIJI (ImageJ, NIH) or Imaris (Bitplane) for analysis. Individual optical planes or max Z-stacks of sequential optical sections were used to assemble figures. Identification of the basal stem cell and differentiated layers/cells was determined with immunofluorescent staining, positional location within the skin, and nuclear morphology. Dermal collagen was capture through second harmonic generation (SHG) of imaging and was used to confirm basal stem cells where immediately adjacent to the basement membrane.

### Chromatin compaction analysis

Data analysis for the chromatin compaction plots was done in Imaris (Bitplane), FIJI (ImageJ), MATLAB, and Prism. We first surfaced the 3D volume of individual nuclei from high-resolution, intravital imaging data. All voxels within the 3D volume were normalized to an 8-bit range of fluorescent intensity inherent to the individual nucleus being surfaced with the top and bottom 0.1% of voxels excluded as outliers. This allowed us to compare chromatin compaction among many different nuclei and among mouse replicates and models despite slight differences in mean fluorescent intensity. Intensity values for each voxel within the 3D volume were binned into 0-256 fluorescent intensity bins, and plotted as a percentage of total nuclear volume to account for differences in nuclear size.

To measure the chromatin compaction within loci of active transcription, we first surfaced individual nuclei and normalized voxel intensity values as described above. We then surfaced the 3D volume of the transcriptionally active locus within each nucleus, applying the nuclear normalization to the voxel intensity values within the transcriptional locus. The intensity values for the voxels within both the nucleus and transcriptional locus were binned and plotted as described above. Coding scripts available upon request.

### Topical drug treatments

To pharmacologically perturb chromatin organization, Trichostatin-A (TSA) was delivered topically to the ear skin. TSA was dissolved in a 10 mg/ml stock solution in dimethyl sulfoxide (DMSO) and then diluted 100X in 100% petroleum jelly (Vaseline; final concentration 1 ug/ml). One hundred micrograms of the TSA/Vaseline mixture was spread evenly on the ear 48 and 24 hours before imaging. A mixture of 100% DMSO in petroleum jelly was used as a vehicle control.

### Statistics and reproducibility

Asterisks denote statistical significance (* *p*<0.05, ** *p*<0.01, *** *p*<0.001 and **** *p*<0.0001). Mean and standard deviation among mice are shown unless otherwise stated. Statistical calculations were performed using the Prism software package (GraphPad, USA).

To test statistical differences between chromatin compaction histograms, we use permutation to test the null hypothesis that the two groups have the same distribution. We define a distance between two groups of histograms. More specifically, we average histogram counts in each group and then calculate the count difference between two groups. Let *H*_*k*_ be the set of all histograms in group *k* and 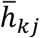 is the average histogram count in the interval *j* of group *k*. Then the distance is defined as

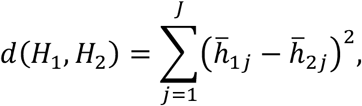

where *J* is the total number of intervals. Based on this definition, we first calculate the distance between the two groups from the observed data. We then perform permutations to derive the null distribution for the distance that there is no group difference. In detail, we permute the labels of the two groups, and calculate the distance for each permuted data set. This is repeated 10,000 times to derive the histogram distance distribution empirically. Lastly, the statistical significance of the observed data is calculated by the proportion of the times that the permuted data lead to a larger distance than that observed. If the p value thus estimated is less than 0.05, we conclude that the histograms of these two groups are significantly different.

### Mouse models

K14-rtTA (Xie et al., 1999), tetO-CDKN1b (Pruitt et al., 2013), tetO-Cre (Strain #:006234), and tdTomato (Madisen et al., 2010) mice were obtained from the Jackson Laboratory. K14H2B-GFP (Tumbar et al., 2004) and K14CreER (Vasioukhin et al., 1999) mice were obtained from Elaine Fuchs, Serum Response Factor (SRF) from Shangqin Guo, K10-rtTA (Muroyama and Lechler, 2017) from Terry Lechler, and MCP-GFP (Lionnet et al., 2011) from Robert Singer. TIGRE-K14-H2B-mCherry mice were generated by the Yale Transgenic Facility. A K14-H2B-mCherry transgene (Mesa et al., 2015) flanked by 2X core sequence of the HS4 chicken beta globin insulator was cloned into a targeting vector (Addgene #92142) that contains homology arms of the mouse TIGRE genomic locus (Madisen et al., 2015). The resulting construct was then used to target into the TIGRE locus via CRISPR/Cas9-mediated genome editing with the gRNA, ACAGAAAACATCCCAAAGTTAGG. One correctly targeted mouse was picked for generating the stable colony.

To block the cell cycle progression of epithelial cells during G1, K14H2B-GFP mice were mated with K14rtTA; tetO-CDKN1b mice and given doxycycline (2mg/ml) in potable water with 2% sucrose. Doxycycline treatment was sustained until imaging was performed the next day. Siblings without the tetO-CDKN1b allele (K14H2B-GFP; K14rtTA) were used as controls. To generate mice deficient for SRF, K14H2B-GFP; K14CreER mice were mated to SRF^fl/wt^; tdTomato^fl/fl^ mice to generate K14H2B-GFP; K14CreER; tdTomato; SRF^fl/wt^ and SRF^wt/wt^ mice in equal proportions. To visualize epidermal cells having initiated transcription of K10, we mated K14H2B-GFP; tetO-Cre mice to K10rtTA; tdTomato mice to yield K14H2B-GFP, K10rtTA; tetO-Cre, tdTomato mice. These mice were given doxycycline (2mg/ml) in potable water with 2% sucrose. Doxycycline treatment was sustained until imaging was performed two days later.

K10MS2 mice were generated by the Yale Transgenic Facility. 24X MS2 repeats (Addgene #31865) were cloned into a vector containing homology arms at the first predicted high-efficiency cut site after the stop codon for the *keratin-10* locus (26bp after stop) (Spille et al., 2019). The resulting construct was then used to target into the keratin-10 locus via CRISPR/Cas9-mediated genome editing with the gRNA, AGTGATCAGGACGATTATTGAGG. Correctly targeted founders were identified for expansion into stable colonies. Mice were born in Mendelian ratios, and heterozygous mice for the K10MS2 allele are phenotypically normal (with normal epidermal structure) which is in agreement with literature of homozygous *keratin-10* knock out mice that exhibit normal differentiation (Reichelt et al., 2001).

Mice from experimental and control groups were randomly selected for either sex for live imaging experiments. All procedures involving animal subjects were performed under the approval of the Institutional Animal Care and Use Committee (IACUC) of the Yale School of Medicine.

### Tamoxifen Induction

To recombine the SRF^fl/wt^/SRF^wt/wt^; K14CreER; tdTomato; K14H2B-GFP mice, we gave a single dose of tamoxifen (20mg/kg body weight in corn oil) by intraperitoneal injection 6 days before the final time point, immediately after imaging the day 0 timepoint.

### PHATE (Potential of Heat-diffusion for Affinity-based Transition Embedding) Analysis and Cell Clustering

MATLAB packages were used for PHATE analysis and Louvain clustering (Blondel et al., 2008; Moon et al., 2019). PHATE was performed on a matrix composed of individual cells and 27 normalized fluorescent intensity bins. The number of diffusion steps was automatically picked and visualized using 2D PHATE embedding. The cells were clustered using the Louvain algorithm on the same matrix, where the algorithm searched for the 50 nearest neighbors based on Euclidean distance. Clusters were visualized on PHATE space.

## Supplementary Figures

**Supplemental Figure 1:**
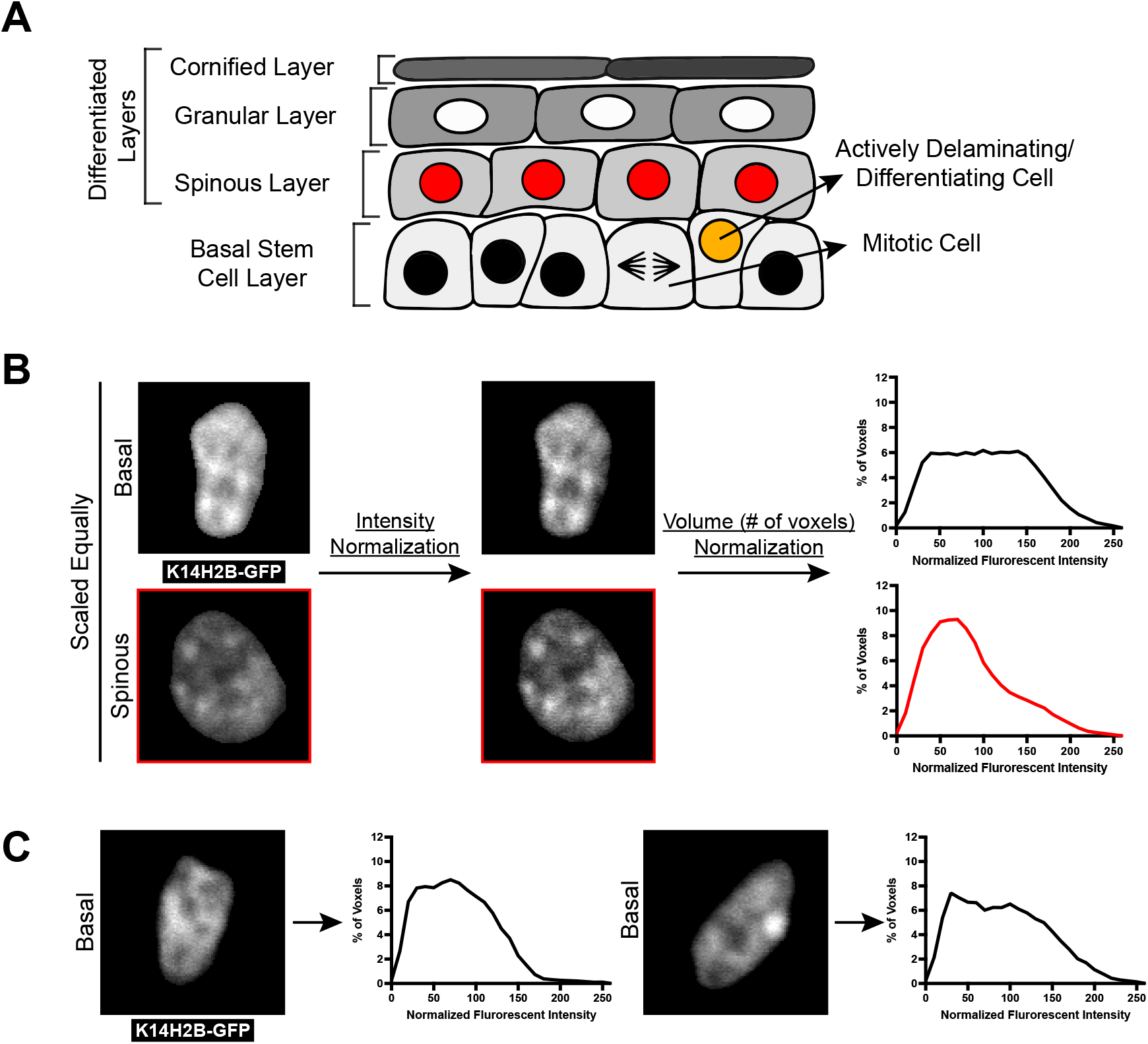
Chromatin compaction state analysis. **(A)** XZ schematic of the mouse epidermis. The basal stem cell layer (bottom) contains actively cycling cells as well as cells that have committed to differentiation and are actively leaving the layer. Cells differentiate apically until they eventually die and form the most outer barrier to the epidermis. **(B)** Quantitative workflow of the chromatin compaction analysis from imaging data (methods). Individual nuclei (left) are surfaced in 3D, exported as raw voxel data, normalized for mean intensity and plotted as a voxel percentage of total nuclear volume. These normalized voxels can then be exported back as scaled, intensity-based images (right). **(C)** Two individual basal nuclei are shown with their corresponding chromatin compaction histograms.

**Supplemental Figure 2:**
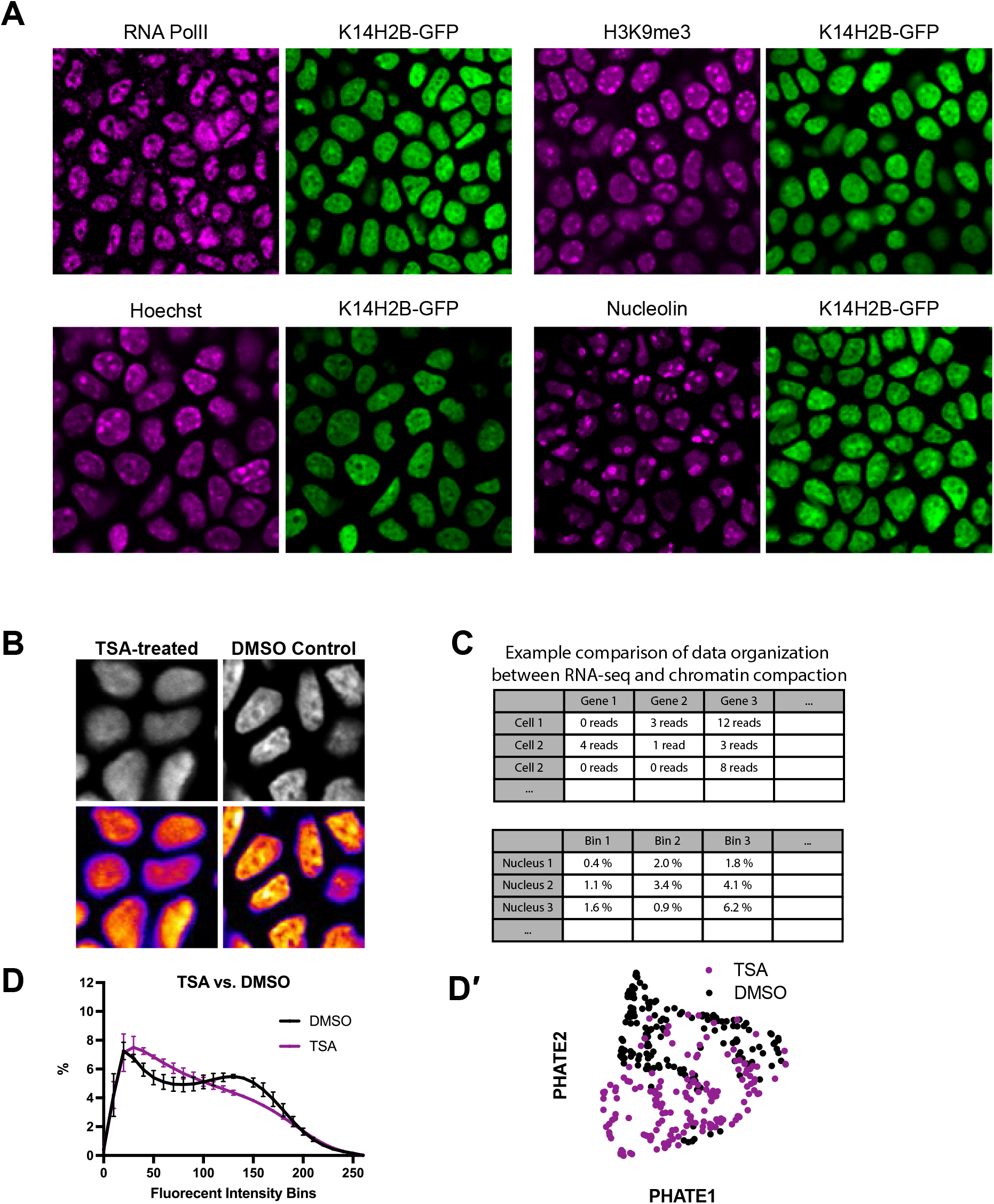
Organization of H2B-GFP allele. **(A)** Representative crops of fixed tissue, single channel stainings of the basal stem cell layer. Genetically encoded K14H2B-GFP signal is shown in green, and immunofluorescent stainings in magenta. **(B)** Representative regions of TSA-treated (left) and DMSO vehicle control (right) mice showing clear disruption of chromatin distribution. K14H2B-GFP intensity seen in greyscale (top) and in the FIRE LUT (bottom). **(C)** Comparison of mock data between scRNA-sequencing data typically applied to the PHATE/Louvain clustering algorithm, and the imaging-based voxel intensity data used. **(D)** Chromatin compaction analysis for TSA-treated and DMSO control nuclei showing altered chromatin compaction state in the TSA-treated mice. N = 150 basal stem cell layer nuclei across 3 mice. **(D′)** PHATE representation of data from (E) showing largely separated clusters of TSA and DMSO-treated mice.

**Supplemental Figure 3:**
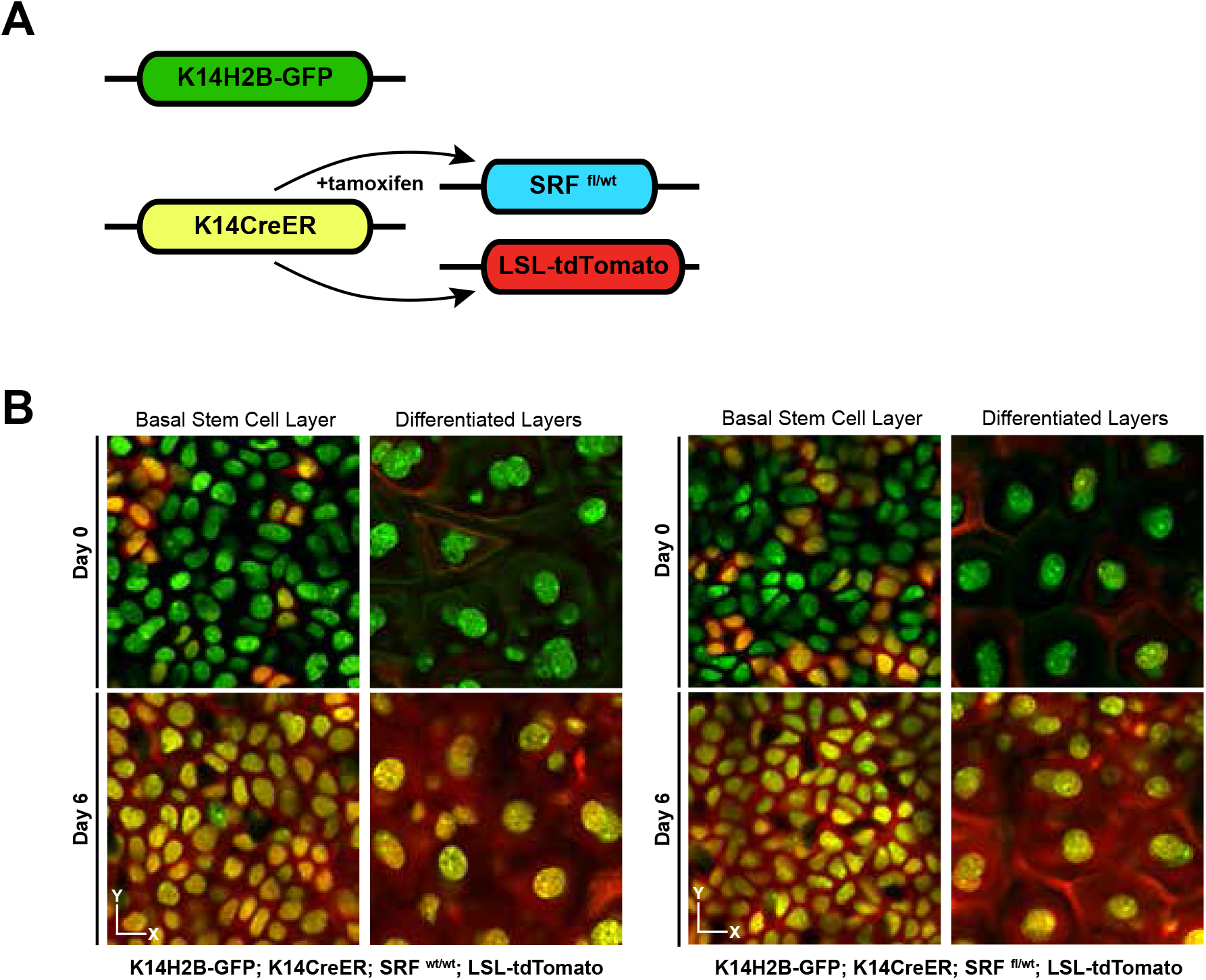
Genetic ablation of SRF alle in the basal stem cell layer. **(A)** Schematic of the inducible genetic system used in figure 2 to recombine and knock down SRF. **(B)** Representative crops from the basal stem cell and differentiated layers on days 0 and 6 in wild-type (top) and SRF heterozygous (bottom) mice. LSL-tdTomato acts as a readout of recombination.

**Supplemental Figure 4:**
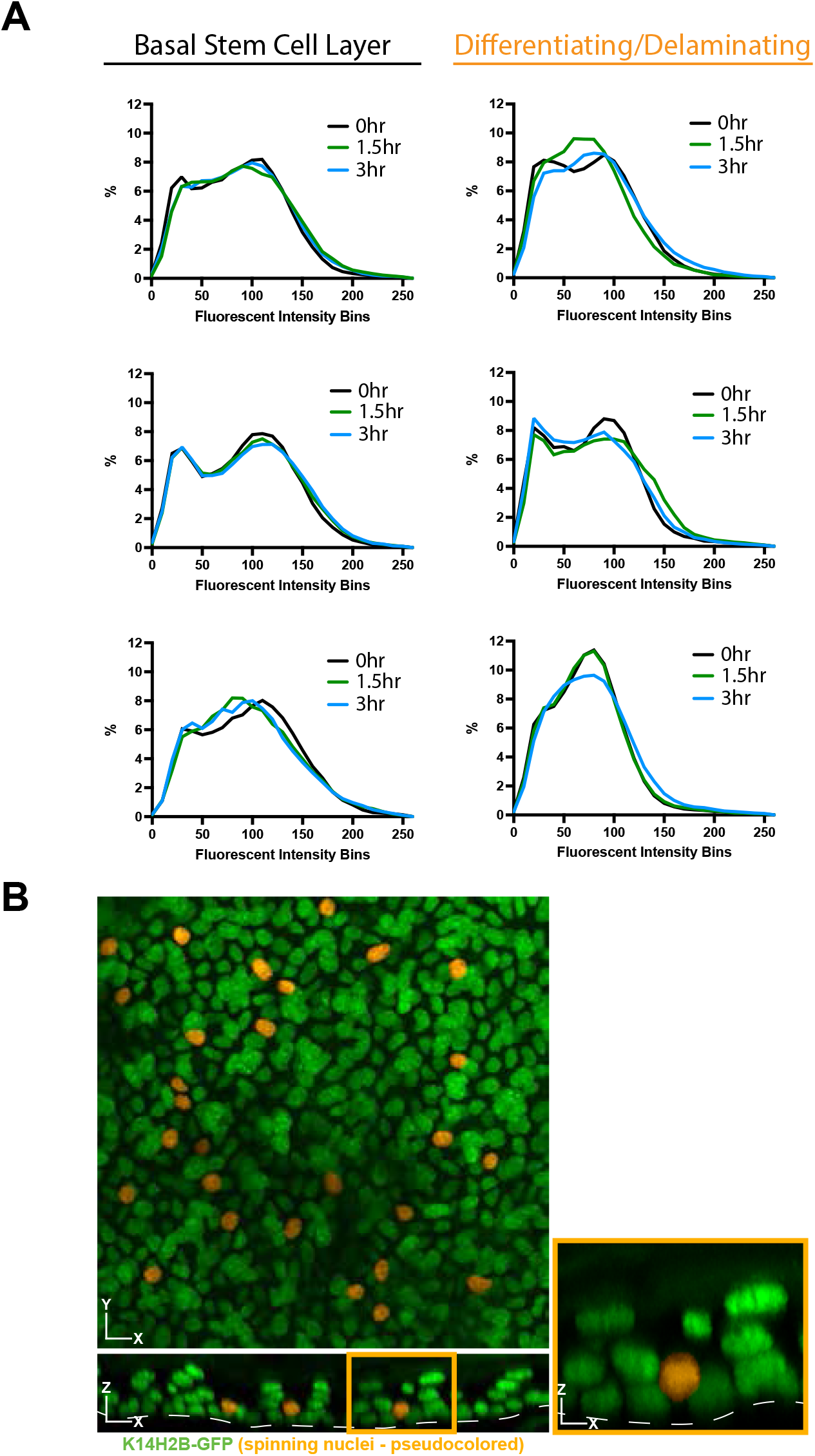
Single cell chromatin compaction dynamics over hours. **(A)** Individual nucleus chromatin compaction profiles from 3 basal stem cell and 3 delaminating/spinning cells at 0-, 1.5-, and 3-hour time points showing little change over this time scale. **(B)** Representative XY (top) and XZ (bottom) max projection FOV of the epidermis (K14H2B-GFP in green) with spinning chromatin cells pseudocolored in orange.

**Supplemental Figure 5:**
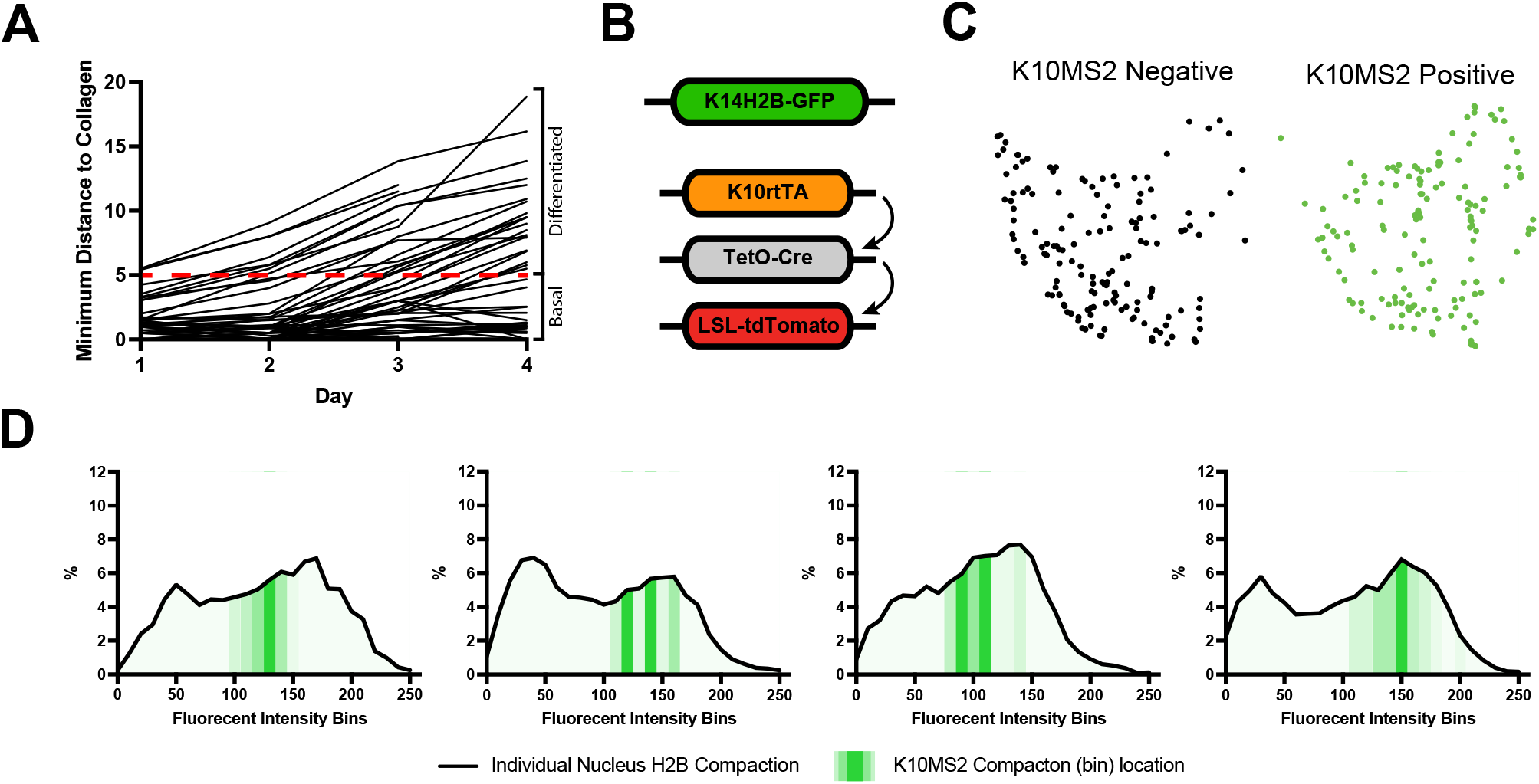
Imaging differentiation-associated cell identity. **(A)** Individual tracking of cells from figure 3D showing trajectory of minimum distance from collagen over 4 days. **(B)** Schematic of the inducible genetic system used in figure 4 to visualize cells within the basal stem cell layer that have started expressing *keratin-10*. **(C)** PHATE representation from figure 5I′ separated by cell status (*keratin-10* “ on” or “ off”). While largely intermixed, there appears a slight preference (though non-significant) for one side of the PHATE cluster vs. the other (“ off” to the left, “ on” to the right). (D) Individual nuclear chromatin compaction profiles for four basal stem cell layer nuclei. K14H2B-mCherry compaction profile shown by black line, and the MCP/MS2 *keratin-10* transcription regions in green translucency.

## Movie Legends

**Movie 1: Chromatin compaction**

A single basal stem cell layer nucleus crop visualizing H2B-GFP intensity. The movie first scans through the greyscale, intensity image (white), then through the FIRE LUT intensities shown in figure 1B, then the three different binned intensities seen in figure 1B and 1C. Nuclear outline is denoted by the white dotted line.

**Movie 2: Spinning chromatin**

An XY field-of-view of the upper, basal stem cell layer in which chromatin (H2B-GFP) can be observed to spin. Timelapse is 3 hours long and looped 3 times.

## References

Amiad-Pavlov, D., Lorber, D., Bajpai, G., Reuveny, A., Roncato, F., Alon, R., Safran, S., and Volk, T. (2021). Live imaging of chromatin distribution reveals novel principles of nuclear architecture and chromatin compartmentalization. Sci. Adv. 7, eabf6251. https://doi.org/10.1126/sciadv.abf6251.

Aragona, M., Sifrim, A., Malfait, M., Song, Y., Van Herck, J., Dekoninck, S., Gargouri, S., Lapouge, G., Swedlund, B., Dubois, C., et al. (2020). Mechanisms of stretch-mediated skin expansion at single-cell resolution. Nature 584, 268–273. https://doi.org/10.1038/s41586-020-2555-7.

Barth, R., Bystricky, K., and Shaban, H.A. (2020). Coupling chromatin structure and dynamics by live super-resolution imaging. Sci. Adv. 6, eaaz2196. https://doi.org/10.1126/sciadv.aaz2196.

Blondel, V.D., Guillaume, J.-L., Lambiotte, R., and Lefebvre, E. (2008). Fast unfolding of communities in large networks. J. Stat. Mech. 2008, P10008. https://doi.org/10.1088/1742-5468/2008/10/P10008.

Bothma, J.P., Garcia, H.G., Esposito, E., Schlissel, G., Gregor, T., and Levine, M. (2014). Dynamic regulation of eve stripe 2 expression reveals transcriptional bursts in living Drosophila embryos. Proc. Natl. Acad. Sci. U.S.A. 111, 10598–10603. https://doi.org/10.1073/pnas.1410022111.

Braun, K.M., Niemann, C., Jensen, U.B., Sundberg, J.P., Silva-Vargas, V., and Watt, F.M. (2003). Manipulation of stem cell proliferation and lineage commitment:visualisation of label-retaining cells in wholemounts of mouse epidermis. Development 130, 5241–5255. https://doi.org/10.1242/dev.00703.

Buenrostro, J.D., Wu, B., Litzenburger, U.M., Ruff, D., Gonzales, M.L., Snyder, M.P., Chang, H.Y., and Greenleaf, W.J. (2015). Single-cell chromatin accessibility reveals principles of regulatory variation. Nature 523, 486–490. https://doi.org/10.1038/nature14590.

Chen, H., Levo, M., Barinov, L., Fujioka, M., Jaynes, J.B., and Gregor, T. (2018). Dynamic interplay between enhancer–promoter topology and gene activity. Nat Genet 50, 1296–1303. https://doi.org/10.1038/s41588-018-0175-z.

Chubb, J.R., Trcek, T., Shenoy, S.M., and Singer, R.H. (2006). Transcriptional Pulsing of a Developmental Gene. Current Biology 16, 1018–1025. https://doi.org/10.1016/j.cub.2006.03.092.

Cockburn, K., Annusver, K., Ganesan, S., Mesa, K.R., Kawaguchi, K., Kasper, M., and Greco, V. (2021). Gradual differentiation uncoupled from cell cycle exit generates heterogeneity in the epidermal stem cell layer. 2021.01.07.425777. https://doi.org/10.1101/2021.01.07.425777.

Dar, R.D., Razooky, B.S., Singh, A., Trimeloni, T.V., McCollum, J.M., Cox, C.D., Simpson, M.L., and Weinberger, L.S. (2012). Transcriptional burst frequency and burst size are equally modulated across the human genome. Proc. Natl. Acad. Sci. U.S.A. 109, 17454–17459. https://doi.org/10.1073/pnas.1213530109.

Doupé, D.P., Klein, A.M., Simons, B.D., and Jones, P.H. (2010). The Ordered Architecture of Murine Ear Epidermis Is Maintained by Progenitor Cells with Random Fate. Developmental Cell 18, 317–323. https://doi.org/10.1016/j.devcel.2009.12.016.

Fan, X., Wang, D., Burgmaier, J.E., Teng, Y., Romano, R.-A., Sinha, S., and Yi, R. (2018). Single Cell and Open Chromatin Analysis Reveals Molecular Origin of Epidermal Cells of the Skin. Developmental Cell 47, 21-37.e5. https://doi.org/10.1016/j.devcel.2018.08.010.

Finn, E.H., Pegoraro, G., Brandão, H.B., Valton, A.-L., Oomen, M.E., Dekker, J., Mirny, L., and Misteli, T. (2019). Extensive Heterogeneity and Intrinsic Variation in Spatial Genome Organization. Cell 176, 1502-1515.e10. https://doi.org/10.1016/j.cell.2019.01.020.

Fritzsch, C., Baumgärtner, S., Kuban, M., Steinshorn, D., Reid, G., and Legewie, S. (2018). Estrogen-dependent control and cell-to-cell variability of transcriptional bursting. Mol Syst Biol 14. https://doi.org/10.15252/msb.20177678.

Gabriele, M., Brandão, H.B., Grosse-Holz, S., Jha, A., Dailey, G.M., Cattoglio, C., Hsieh, T.-H.S., Mirny, L., Zechner, C., and Hansen, A.S. (2022). Dynamics of CTCF-and cohesin-mediated chromatin looping revealed by live-cell imaging. Science 376, 496–501. https://doi.org/10.1126/science.abn6583.

Gdula, M.R., Poterlowicz, K., Mardaryev, A.N., Sharov, A.A., Peng, Y., Fessing, M.Y., and Botchkarev, V.A. (2013). Remodeling of Three-Dimensional Organization of the Nucleus during Terminal Keratinocyte Differentiation in the Epidermis. Journal of Investigative Dermatology 133, 2191–2201. https://doi.org/10.1038/jid.2013.66.

Golkaram, M., Jang, J., Hellander, S., Kosik, K.S., and Petzold, L.R. (2017). The Role of Chromatin Density in Cell Population Heterogeneity during Stem Cell Differentiation. Sci Rep 7, 13307. https://doi.org/10.1038/s41598-017-13731-3.

Hiratsuka, T., Fujita, Y., Naoki, H., Aoki, K., Kamioka, Y., and Matsuda, M. (2015). Intercellular propagation of extracellular signal-regulated kinase activation revealed by in vivo imaging of mouse skin. ELife 4, e05178. https://doi.org/10.7554/eLife.05178.

Iida, S., Shinkai, S., Itoh, Y., Tamura, S., Kanemaki, M.T., Onami, S., and Maeshima, K. (2022). Single-nucleosome imaging reveals steady-state motion of interphase chromatin in living human cells. Sci. Adv. 8, eabn5626. https://doi.org/10.1126/sciadv.abn5626.

Jin, W., Tang, Q., Wan, M., Cui, K., Zhang, Y., Ren, G., Ni, B., Sklar, J., Przytycka, T.M., Childs, R., et al. (2015). Genome-wide detection of DNase I hypersensitive sites in single cells and FFPE tissue samples. Nature 528, 142–146. https://doi.org/10.1038/nature15740.

Kanda, T., Sullivan, K.F., and Wahl, G.M. (1998). Histone–GFP fusion protein enables sensitive analysis of chromosome dynamics in living mammalian cells. Current Biology 8, 377–385. https://doi.org/10.1016/S0960-9822(98)70156-3.

Kurimoto, K., Yabuta, Y., Hayashi, K., Ohta, H., Kiyonari, H., Mitani, T., Moritoki, Y., Kohri, K., Kimura, H., Yamamoto, T., et al. (2015). Quantitative Dynamics of Chromatin Remodeling during Germ Cell Specification from Mouse Embryonic Stem Cells. Cell Stem Cell 16, 517–532. https://doi.org/10.1016/j.stem.2015.03.002.

Lai, B., Gao, W., Cui, K., Xie, W., Tang, Q., Jin, W., Hu, G., Ni, B., and Zhao, K. (2018). Principles of nucleosome organization revealed by single-cell micrococcal nuclease sequencing. Nature 562, 281–285. https://doi.org/10.1038/s41586-018-0567-3.

Li, D., Liu, J., Yang, X., Zhou, C., Guo, J., Wu, C., Qin, Y., Guo, L., He, J., Yu, S., et al. (2017). Chromatin Accessibility Dynamics during iPSC Reprogramming. Cell Stem Cell 21, 819-833.e6. https://doi.org/10.1016/j.stem.2017.10.012.

Li, Y., Eshein, A., Virk, R.K.A., Eid, A., Wu, W., Frederick, J., VanDerway, D., Gladstein, S., Huang, K., Shim, A.R., et al. (2021). Nanoscale chromatin imaging and analysis platform bridges 4D chromatin organization with molecular function. Sci. Adv. 7, eabe4310. https://doi.org/10.1126/sciadv.abe4310.

Lin, C., Hindes, A., Burns, C.J., Koppel, A.C., Kiss, A., Yin, Y., Ma, L., Blumenberg, M., Khnykin, D., Jahnsen, F.L., et al. (2013). Serum response factor controls transcriptional network regulating epidermal function and hair follicle morphogenesis. J Invest Dermatol 133, 608–617. https://doi.org/10.1038/jid.2012.378.

Lionnet, T., Czaplinski, K., Darzacq, X., Shav-Tal, Y., Wells, A.L., Chao, J.A., Park, H.Y., de Turris, V., Lopez-Jones, M., and Singer, R.H. (2011). A transgenic mouse for in vivo detection of endogenous labeled mRNA. Nat Methods 8, 165–170. https://doi.org/10.1038/nmeth.1551.

Madisen, L., Zwingman, T.A., Sunkin, S.M., Oh, S.W., Zariwala, H.A., Gu, H., Ng, L.L., Palmiter, R.D., Hawrylycz, M.J., Jones, A.R., et al. (2010). A robust and high-throughput Cre reporting and characterization system for the whole mouse brain. Nat Neurosci 13, 133–140. https://doi.org/10.1038/nn.2467.

Madisen, L., Garner, A.R., Shimaoka, D., Chuong, A.S., Klapoetke, N.C., Li, L., van der Bourg, A., Niino, Y., Egolf, L., Monetti, C., et al. (2015). Transgenic Mice for Intersectional Targeting of Neural Sensors and Effectors with High Specificity and Performance. Neuron 85, 942–958. https://doi.org/10.1016/j.neuron.2015.02.022.

Mardaryev, A.N., Gdula, M.R., Yarker, J.L., Emelianov, V.U., Poterlowicz, K., Sharov, A.A., Sharova, T.Y., Scarpa, J.A., Joffe, B., Solovei, I., et al. (2014). p63 and Brg1 control developmentally regulated higher-order chromatin remodelling at the epidermal differentiation complex locus in epidermal progenitor cells. Development 141, 3437–3437. https://doi.org/10.1242/dev.115725.

Mesa, K.R., Rompolas, P., Zito, G., Myung, P., Sun, T.Y., Brown, S., Gonzalez, D.G., Blagoev, K.B., Haberman, A.M., and Greco, V. (2015). Niche-induced cell death and epithelial phagocytosis regulate hair follicle stem cell pool. Nature 522, 94–97. https://doi.org/10.1038/nature14306.

Mesa, K.R., Kawaguchi, K., Cockburn, K., Gonzalez, D., Boucher, J., Xin, T., Klein, A.M., and Greco, V. (2018). Homeostatic Epidermal Stem Cell Self-Renewal Is Driven by Local Differentiation. Cell Stem Cell 23, 677-686.e4. https://doi.org/10.1016/j.stem.2018.09.005.

Moon, K.R., van Dijk, D., Wang, Z., Gigante, S., Burkhardt, D.B., Chen, W.S., Yim, K., Elzen, A. van den, Hirn, M.J., Coifman, R.R., et al. (2019). Visualizing structure and transitions in high-dimensional biological data. Nat Biotechnol 37, 1482–1492. https://doi.org/10.1038/s41587-019-0336-3.

Muroyama, A., and Lechler, T. (2017). A transgenic toolkit for visualizing and perturbing microtubules reveals unexpected functions in the epidermis. ELife 6, e29834. https://doi.org/10.7554/eLife.29834.

Oudelaar, A.M., Beagrie, R.A., Gosden, M., de Ornellas, S., Georgiades, E., Kerry, J., Hidalgo, D., Carrelha, J., Shivalingam, A., El-Sagheer, A.H., et al. (2020). Dynamics of the 4D genome during in vivo lineage specification and differentiation. Nat Commun 11, 2722. https://doi.org/10.1038/s41467-020-16598-7.

Patel, A.P., Tirosh, I., Trombetta, J.J., Shalek, A.K., Gillespie, S.M., Wakimoto, H., Cahill, D.P., Nahed, B.V., Curry, W.T., Martuza, R.L., et al. (2014). Single-cell RNA-seq highlights intratumoral heterogeneity in primary glioblastoma. Science 344, 1396–1401. https://doi.org/10.1126/science.1254257.

Paulsen, J., Liyakat Ali, T.M., Nekrasov, M., Delbarre, E., Baudement, M.-O., Kurscheid, S., Tremethick, D., and Collas, P. (2019). Long-range interactions between topologically associating domains shape the four-dimensional genome during differentiation. Nat Genet 51, 835–843. https://doi.org/10.1038/s41588-019-0392-0.

Pelham-Webb, B., Murphy, D., and Apostolou, E. (2020). Dynamic 3D Chromatin Reorganization during Establishment and Maintenance of Pluripotency. Stem Cell Reports 15, 1176–1195. https://doi.org/10.1016/j.stemcr.2020.10.012.

Pruitt, S.C., Freeland, A., Rusiniak, M.E., Kunnev, D., and Cady, G.K. (2013). Cdkn1b overexpression in adult mice alters the balance between genome and tissue ageing. Nat Commun 4, 2626. https://doi.org/10.1038/ncomms3626.

Reichelt, J., Büssow, H., Grund, C., and Magin, T.M. (2001). Formation of a normal epidermis supported by increased stability of keratins 5 and 14 in keratin 10 null mice. Mol Biol Cell 12, 1557–1568. https://doi.org/10.1091/mbc.12.6.1557.

Schweizer, J., Kinjo, M., Fürstenberger, G., and Winter, H. (1984). Sequential expression of mRNA-encoded keratin sets in neonatal mouse epidermis: Basal cells with properties of terminally differentiating cells. Cell 37, 159–170. https://doi.org/10.1016/0092-8674(84)90311-8.

Shue, Y.T., Lee, K.T., Walters, B.W., Ong, H.B., Silvaraju, S., Lam, W.J., and Lim, C.Y. (2020). Dynamic shifts in chromatin states differentially mark the proliferative basal cells and terminally differentiated cells of the developing epidermis. Epigenetics 15, 932–948. https://doi.org/10.1080/15592294.2020.1738028.

Spille, J.-H., Hecht, M., Grube, V., Cho, W., Lee, C., and Cissé, I.I. (2019). A CRISPR/Cas9 platform for MS2-labelling of single mRNA in live stem cells. Methods 153, 35–45. https://doi.org/10.1016/j.ymeth.2018.09.004.

Suter, D.M., Molina, N., Gatfield, D., Schneider, K., Schibler, U., and Naef, F. (2011). Mammalian Genes Are Transcribed with Widely Different Bursting Kinetics. Science 332, 472–474. https://doi.org/10.1126/science.1198817.

Tumbar, T., Guasch, G., Greco, V., Blanpain, C., Lowry, W.E., Rendl, M., and Fuchs, E. (2004). Defining the epithelial stem cell niche in skin. Science 303, 359–363. https://doi.org/10.1126/science.1092436.

Vasioukhin, V., Degenstein, L., Wise, B., and Fuchs, E. (1999). The magical touch: Genome targeting in epidermal stem cells induced by tamoxifen application to mouse skin. Proc. Natl. Acad. Sci. U.S.A. 96, 8551–8556. https://doi.org/10.1073/pnas.96.15.8551.

Verdoni, A.M., Ikeda, S., and Ikeda, A. (2010). Serum response factor is essential for the proper development of skin epithelium. Mamm Genome 21, 64–76. https://doi.org/10.1007/s00335-009-9245-y.

Wang, S., Drummond, M.L., Guerrero-Juarez, C.F., Tarapore, E., MacLean, A.L., Stabell, A.R., Wu, S.C., Gutierrez, G., That, B.T., Benavente, C.A., et al. (2020). Single cell transcriptomics of human epidermis identifies basal stem cell transition states. Nat Commun 11, 4239. https://doi.org/10.1038/s41467-020-18075-7.

Xie, W., Chow, L.T., Paterson, A.J., Chin, E., and Kudlow, J.E. (1999). Conditional expression of the ErbB2 oncogene elicits reversible hyperplasia in stratified epithelia and up-regulation of TGFα expression in transgenic mice. Oncogene 18, 3593–3607. https://doi.org/10.1038/sj.onc.1202673.

